# CRB2 Depletion Induces YAP Signaling and Disrupts Mechanosensing in Podocytes

**DOI:** 10.1101/2024.10.22.619513

**Authors:** Yingyu Sun, Nils M. Kronenberg, Sidharth K. Sethi, Surjya N. Dash, Maria E. Kovalik, Benjamin Sempowski, Shelby Strickland, Rupresh Raina, C. John Sperati, Xuefei Tian, Shuta Ishibe, Gentzon Hall, Malte C. Gather

## Abstract

Focal Segmental Glomerulosclerosis (FSGS) is a histologic lesion caused by a variety of injurious stimuli that lead to dysfunction/loss of glomerular visceral epithelial cells (i.e. podocytes). Pathogenic mutations in CRB2, encoding the type 1 transmembrane protein Crumb 2 Homolog Protein, have been shown to cause early-onset corticosteroid-resistant nephrotic syndrome (SRNS)/FSGS. Here, we identified a 2-generation East Asian kindred (DUK40595) with biopsy-proven SRNS/FSGS caused by a compound heterozygous mutation in CRB2 comprised of the previously described truncating mutation p.Gly1036_Alafs*43 and a rare 9-bp deletion mutation p.Leu1074_Asp1076del. Because compound heterozygous mutations involving the truncating p.Gly1036_Alafs*43 variant have been associated with reduced CRB2 expression in podocytes and autosomal recessive SRNS/FSGS, we sought to define the pathogenic effects of CRB2 deficiency in podocytes. We show that CRB2 knockdown induces YAP activity and target gene expression in podocytes. It upregulates YAP-mediated mechanosignaling and increases the density of focal adhesion and F-actin. Using Elastic Resonator Interference Stress Microscopy (ERISM), we demonstrate that CRB2 knockdown also enhances podocyte contractility in a substrate stiffness-dependent manner. The knockdown effect decreases with increasing substrate stiffness, indicating impaired mechanosensing in CRB2 knockdown cells at low substrate stiffness. While the mechanical activation of CRB2 knockdown cells is associated with increased YAP activity, the enhanced cell contractility is not significantly reduced by the selective YAP inhibitors K-975 and verteporfin, suggesting that multiple pathways may be involved in mechanosignaling downstream of CRB2. Taken together, these studies provide the first evidence that CRB2 deficiency may impair podocyte mechanotransduction via disruption of YAP signaling in podocytes.

## INTRODUCTION

Focal Segmental Glomerulosclerosis (FSGS) is a histologic injury caused by dysfunction or loss of glomerular visceral epithelial cells (i.e. podocytes) (1). In over 70% of patients with FSGS, the most common clinical manifestation is nephrotic syndrome (NS) and nearly 50% of patients with FSGS progress to end-stage kidney disease (ESKD) over a decade (2). It is estimated that approximately 20% of patients receiving hemodialysis have biopsy-proven FSGS (3–5). Currently, there are no targeted treatments for the lesion and standard-of-care therapies focus on controlling hypertension, proteinuria reduction, and blockade of the renin angiotensin system (6). Immunosuppressive therapies may also be added in some cases based on the observation that immune system dysregulation can contribute to podocyte injury (7). The clinical and economic burden of FSGS care is substantial (8) as the lesion is estimated to cause disease in nearly 40,000 patients and has an incidence of approximately 5,000 cases per year in the United States (US) (3, 6, 9). Notably, the incidence of FSGS has increased more than 10-fold over the past three decades and it is now the most common primary glomerular lesion that causes ESKD in the US (7, 10). Worldwide, the estimated incidence of FSGS ranges from 1.2 - 21 cases per million population with 7 - 10% of cases occurring in children and 20 - 40% occurring in adults with NS (11).

FSGS can result from a variety of stimuli, contributing to a heterogeneity of the histologic lesion and variability in the clinical presentation and treatment response (1). Regardless of the cause, it is clear that the final common pathway to developing FSGS involves podocyte injury or loss (7, 12–14). Podocytes are an essential cellular component of the tripartite glomerular filtration barrier (GFB) as they synthesize and maintain the filtration slit diaphragm (SD) (15, 16). The SD is a “ladder-like” assembly of proteins that bridge the interdigital spaces between adjacent podocyte foot processes (17–19). It is thought that this dynamic, macromolecular assembly functions like a sieve to provide size and charge selectivity for ultrafiltration (17–19). To date, mutations in over 60 podocyte-expressed genes have been shown to cause familial SRNS/FSGS (20–22). Mutations in the gene that encodes Crumbs Homolog 2 (*CRB2*; FSGS9) have been shown to cause early-onset, steroid-resistant nephrotic syndrome (SRNS)/FSGS (23–32). CRB2 is an essential SD protein that organizes in clusters in the podocyte apical membrane and interacts with nephrin and other SD proteins (Figure S1) (26). CRB2 is one of three members of the Crumbs family (CRB1, CRB2 and CRB3) of apical polarity-related type I transmembrane proteins (33). Crumbs proteins are widely expressed in epithelial cells and regulate cell shape, polarity and intracellular signaling (33). While the expression of CRB1 is largely restricted to the central nervous system, CRB2 and CRB3 are both expressed throughout the body, including in the kidneys (34, 35). Notably, inherited glomerular disease has only been associated with *CRB2* mutations and no known human genetic disease has been associated with mutations in *CRB3* (31, 34, 36). Here we report the identification of a rare compound heterozygous *CRB2* mutation in a 2-generation East Asian kindred with biopsy-proven FSGS. Affected family members expressed the previously reported truncating frameshift mutation p.Gly1036AlafsTer43 (23) and a rare, in-frame 9-bp deletion mutation, p.Leu1074_Asp1076del. We hypothesized that these mutations would exert a loss-of-function effect and sought to explore the pathogenic mechanisms of the compound heterozygous mutation in podocytes using an *in vitro* model of CRB2 deficiency combined with extensive measurements of the mechanical activity of podocytes.

Despite a growing body of evidence demonstrating the essential role of CRB2 in podocyte function and viability, few mechanistic insights into the pathobiology of CRB2 deficiency have been defined. To date, truncating *CRB2* mutations have been associated with reduced SD protein expression, increased apoptotic signaling, impaired adhesion, and dysregulated sphingosine 1-phosphate receptor (S1PR1) expression/phosphorylation in podocytes (23, 27, 32). CRB2 is widely recognized as a critical upstream regulator of Hippo pathway signaling in epithelial cells and dysregulated Hippo signaling in podocytes has been implicated in the pathogenesis of glomerular disease (37–39). Several studies have identified the Yes-associated Protein (YAP) as a critical downstream mediator of Hippo signaling in podocyte health and disease (40). YAP is an inducible transcriptional co-activator that has been shown to regulate podocyte apoptosis, actin cytoskeletal dynamics, adhesion and mechanotransduction (40–42). While the role of YAP in these processes is well established in podocytes, the influence of CRB2 on YAP activity remains unclear.

In the present study, we sought to clarify the role of CRB2 as an upstream regulator of YAP mechanosignaling in podocytes. Using siRNA-mediated *CRB2* knockdown in conditionally immortalized human podocytes, we demonstrate that YAP target gene and protein expression, focal adhesion density and mechanosignaling are significantly increased. Using Elastic Resonator Interference Stress Microscopy (ERISM), a recently developed force microscopy modality for long-term mapping of mechanical activity of cells (43, 44), we demonstrate that contractile force development is significantly increased in CRB2 knockdown podocytes. In addition, CRB2 knockdown podocytes exhibit a higher F-actin density and paxillin expression and phosphorylation while their cell area is significantly decreased. In order to examine if CRB2 knockdown has an impact on how podocytes adjust contractile forces to the mechanical properties of the micro-environment (mechanosensing), we performed a detailed characterization of podocyte force development over a range of substrate stiffnesses throughout differentiation. In line with the proposed function of CRB2 being an upstream regulator of YAP activation, we found that the relative effect of CRB2 knockdown on cell contractility decreases with substrate stiffness. Interestingly, treatment of CRB2 knockdown podocytes with the YAP inhibitors verteporfin and K-975 did not reduce podocyte contractile force development relative to controls, suggesting that other pathways than YAP signaling may also contribute to mechanosignaling downstream of CRB2. Taken together, these findings provide the first evidence that CRB2 deficiency may impair podocyte mechanotransduction via disruption of YAP signaling in podocytes.

## MATERIALS AND METHODS

### Genome sequencing

Proband blood DNA was subjected to targeted gene capture and sequencing to a mean > 80-100X coverage on the Illumina sequencing platform through MedGenome Labs Pvt. Ltd, Bangalore, India. The sequences obtained were aligned to the human reference genome (GRCh37/hg19) using the Burrows-Wheeler Alignment (BWA) program (45, 46) and were analyzed using Picard and the Genome Analysis Toolkit (GATK) version 3.6 (47, 48) to identify variants relevant to the clinical indication. Gene annotation of the variants was performed using the Variant Effect Predictor program (49) against the Ensembl release 87 human gene model (http://www.ensembl.org/). Clinically relevant mutations were annotated using published variants in literature and the ClinVaar, OMIM, GWAS, HGMD and SwissVar disease databases (50–54). Common variants were filtered based on allele frequency in 1000 Genome Phase 3, ExAC, EVS, dbSNP147, 1000 Japanese Genome and regional Indian population database (55–57). Only non-synonymous and splice-variants found in the exome panel consisting of 8,322 genes were used for clinical interpretation.

### *CRB2* Primers and Direct Sequencing

*CRB2* Family PCR

*CRB2*_Nested2_F: AGATTCTGCCAGGAGTTGCT

*CRB2*_Nested2_R: AACACAACGCCTGGACATTC

Chr19:123372844+123374338 1495bp

PCR was performed on previously extracted samples from blood using the PrimeStar GXL Polymerase kit, with final concentrations of 1x PrimerStar GXL Buffer, 200 µM each dNTP, 0.1µM each primer, 1.25 U/50ml PrimeStar GXL Polymerase, and 0.5 M betaine. PCR amplification was performed using a thermocycler (GeneAmp 9700) with 30 cycles of 98 °C for 10 s, 67 °C for 15 s, and 68 °C for 15 s using the following primers, *CRB2*_Nested2_F: AGATTCTGCCAGGAGTTGCT and *CRB2*_Nested2_R: AACACAACGCCTGGACATTC. Products were then confirmed via a 1% agarose gel run at 100 V for 30 min and Sanger Sequenced using both forward and reverse primers at GeneWiz. Sequencing files were uploaded to Sequencher, trimmed, and assembled to the reference sequence. Heterozygous bases were called using the internal software, and all mismatches to the reference sequence were visually checked for accuracy using the chromatogram sequences.

### Protein structure modelling

In silico modeling of the CRB2 p.Leu1074_Asp1076del variant was performed using the I-TASSER protein structure prediction software (http://zhanglab.ccmb.med.umich.edu/I-TASSER/) (58, 59). Structural renderings were generated using the PyMOL molecular graphics program.

### Lentiviral Vector production and transduction of siScr and si*CRB2* Lines

Lentiviral vectors were generated using the transient infection protocol described in (60). Briefly, 15 μg of vector plasmid, 10 μg of psPAX2 packaging plasmid (Addgene #12260 generated in Dr Didier Trono’s lab, EPFL, Switzerland), 5 μg of pMD2.G envelope plasmid (Addgene #12259, generated in Dr Trono’s lab) and 2.5 μg of pRSV-Rev plasmid (Addgene #12253, generated in Dr Trono’s lab) were transfected into 293 T cells. Vector particles were collected from the filtered conditioned medium at 72 h post-transfection. The particles were purified using the sucrose-gradient method and concentrated > 250-fold by ultracentrifugation (2 h at 20 000 rpm). Vector and viral stocks were aliquoted and stored at −80 °C. Conditionally immortalized podocytes were plated on collagen I-coated 6-well flasks and maintained under growth-permissive conditions until 70% confluence was achieved. Cells were then transduced with 4 µL of siScr and si*CRB2* lentiviruses at a concentration of 1.0 x 10^5^ particles/µL in serum-free media. Cells were incubated overnight before washing with complete media and further incubation for 48 hours. At confluence, antibiotic selection was initiated for 5-7 days. Surviving cells were expanded in complete media prior to transition to growth-restrictive conditions at confluence. After 12 days of differentiation, cells were harvested and assayed for *CRB2* mRNA expression as described below.

### Molecular cloning of luciferase reporter lines

The 8x GTIIC-luciferase plasmid was obtained from Addgene (#34615; gift of Dr. Stefano Piccolo). The following primers were used to amplify the 8x GTIIC-luciferase region: Forward-CCGACTCGAGAATTACACGGC; Reverse-CTATCGATATCTACCGAGCTCTTACGC. The XhoI and EcoRV restriction sites were added at the 5’ and 3’ ends, respectively. The fragment was cloned into a standard SIN-LTR lentiviral vector cassette. Plasmid #1773 (Addgene, gift of Dr. Bob Weinberg) was then amplified using the following primers: Forward-TGGACTCGAGTCGACCCTGT and Reverse-GATCCCGCTCGAGATCTACTCTATTCC. The fragment carrying SV40-promoter Hygromycin resistant gene flanked by XhoI restriction sites was cloned into an intermediate vector to create the final pBK909 plasmid used for cell transfection. All vectors were validated by direct Sanger sequencing and by digestion. The final plasmid was amplified and prepared for transfection as described in (61). Once siScr and si*CRB2* lines were established, cells were plated on collagen I-coated 6-well flasks and maintained under growth-permissive conditions until 70% confluence was achieved. Cells were then transduced with 4 µl of the 8x GTIIC lentivirus at a concentration of 1.0 x 10^5^ particles/µl in serum-free media. Cells were then incubated overnight before washing with complete media and further incubation for 48 hours. At confluence, antibiotic selection was initiated for 5-7 days. Surviving cells were expanded in complete media.

### Cell culture

Human conditionally immortalized podocytes contained human telomerase reverse transcriptase (hTERT). The thermosensitive SV40 large T antigen was kindly provided by J. Kopp (Bethesda, MD, USA). Both siScr and si*CRB2* podocytes were established through lentiviral transduction with scrambled and *CRB2* siRNA (Applied Biological Materials), respectively. Podocytes were incubated on collagen-I-coated flasks (Corning) under permissive temperature (33 °C) and 5% CO_2_ and in RPMI 1640 (R8758, Gibco) supplemented with 10% of fetal bovine serum (FBS) (A5209402, Gibco), 1% of penicillin-streptomycin (15140122, Gibco) and 1% of insulin-transferrin-selenium (ITS) (354351, Corning). Podocytes were split at a confluency around 90%. Both cell lines were seeded onto an ERISM substrate before performing the thermoswitch to the differentiation conditions at 37 °C, 5% CO_2_ for 12 days. Culture medium was refreshed every 2 to 3 days. Podocytes were exposed to the YAP-transcriptional inhibitors verteporfin (Sigma-Aldrich) and K-975 (MedChemExpress) and to the mTOR inhibitor rapamycin (MedChemExpress) for 24 hours using the concentrations indicated in Figure 7. All groups of the inhibitor study were treated with 0.2% (v/v) of DMSO.

### RNA Extraction and qPCR

Cells were grown to 95% confluency in a T75 collagen coated flask, rinsed with 1xPBS, and incubated for 5 min at room temperature in 5 ml of trypsin. Media was added to each sample to neutralize the trypsin, and all flask contents were transferred to a 15 ml conical tube. The conicals were centrifuged at 1500 RPM for 5 min with 2 washes of 1xPBS. RNA was extracted using the Qiagen RNeasy miniprep kit according to manufacturer’s instructions. RNA eluates were quantified using Nanodrop and normalized to 500 ng/µl. cDNA was generated using the BioRad iScript cDNA synthesis kit in 2 μg of RNA input in a 40 μl reaction volume according to manufacturer’s instructions. Taqman Gene Expression Assays (Invitrogen) were used in conjunction with Taqman Fast Advanced Master Mix (Invitrogen) using 0.5 μl of Assay mix, 5 μl of Master Mix, 3.5μl of nuclease-free water, and 1µl of cDNA per reaction. The assays were run on a ViiA7 with manufacturer’s cycling parameters. All samples were run in triplicate along with no-template controls. β-Actin was used as the internal control gene and the relative quantification was calculated using the ViiA7 software.

### Immunohistochemistry

The immunofluorescence staining of human kidney biopsy specimen was performed as previously described (62). Briefly, 4 µm sections of paraffin-embedded human kidney tissues were deparaffinized with xylene, followed by hydration using a gradient of ethanol concentrations at room temperature. Antigen epitope retrieval was performed by incubating the kidney sections in 10 mM sodium citrate buffer with 0.05% Tween 20 (pH = 6.0) for 10 min. Subsequently, the sections were blocked with 3% BSA in 1x PBS buffer for 1 hour at room temperature. The sections were then incubated overnight at 4 °C with *CRB2* rabbit polyclonal antibody (1:100, VWR, AP5724B) and nephrin guinea pig polyclonal antibody (1:200, Progen, GP-N2). Following the incubation, the slides were washed three times with 1x PBS and incubated with Alexa Fluor 488 goat anti-rabbit antibody (1:200, Invitrogen, A11034) and Alexa Fluor 594 goat anti-guinea pig antibody (1:200, Invitrogen, A11076) at room temperature for 1 hour in the dark. After three additional washes with 1x PBS, the sections were mounted with DAPI Slowfade (Invitrogen, S36938). Images were acquired using an Andor CSU-WDi spinning disk confocal microscope equipped with a Nikon Ti-E CFI Plan Apochromat Lambda 60× oil immersion objective.

### Immunocytochemistry

Podocytes were seeded either in petri dishes (80416, Ibidi) for nephrin and synaptopodin, on glass coverslips for FAK1, CRB2 and TRPC6, or on ERISM substrates for paxillin immunocytochemistry. The latter was combined with ERISM measurements; for this, a coordinate array of 5 cells per chamber were recorded before taking a full-range ERISM scan of the array to yield the reference local thickness of the field of view and further fast one-minimum ERISM scans (63) were performed prior to the fixation (see below).

Cells were fixed with 4% paraformaldehyde (PFA) at room temperature for 15 min (petri dishes and ERISM substrates), or 20 min (glass coverslips), and were washed three times in 1x PBS with 5 min of incubation. Except for membrane staining, cells were permeabilized using 0.1% or 0.2% Triton X-100 in 1x PBST (1xPBS + 0.2% Tween 20) for 5 min, after which cells were blocked with 1% or 2% BSA for 30 min and then rinsed with 1x PBST. After washing the cells for 5 min twice, they were incubated with the following primary antibodies at 4°C overnight: nephrin rabbit polyclonal antibody (1:200, Thermo Fisher Scientific, PA5-20330), synaptopodin mouse monoclonal antibody (1:200, Progen Biotechnik, 61094), FAK1mouse monoclonal antibody (1:100, Millipore Sigma, 05-537), CRB2 rabbit polyclonal antibody (1:100, BIOSS, BS-14046R), TRPC6 mouse monoclonal antibody (1:100, Abcam, 105845), and at room temperature for one hour: paxillin rabbit monoclonal antibody (1:1200, Cell Signaling, 50195) in 1% BSA. After washing 3 to 5 times with 1X PBS or 1X PBST, cells were incubated in the following secondary antibodies for one hour at room temperature in dark: Alexa Fluor Plus 647 donkey anti-mouse (1:800) (Thermo Fisher, A32787), Cy5 donkey anti-rabbit (1:400, Jackson ImmunoResearch, 711-175-152), Alexa Fluor 555/488 donkey anti-mouse and donkey anti-rabbit (1:1000, ThermoFisher, A-31570 and A-21206), TRITC-phalloidin (1:500, Sigma Aldrich, FAK100), Alexa Fluor 488/555-phalloidin (1:1000, ThermoFisher, A12379 and A30106) and DAPI (1:800 or 1:2000). The stained cells were washed 3 to 5 times in 1xPBS or 1xPBST. Cells on glass coverslips were placed on ProLongTM Gold Antifade Mountant (ThermoFisher); the other samples were imaged in 1x PBS.

For nephrin/ or synaptopodin/DAPI staining in petri dishes, cells were imaged under a Leica Stellaris 8 confocal microscope at HCNB; for phalloidin/CRB2/TRPC6 and FAK1/DAPI on glass coverslips a Leica SP8-confocal microscope at the Duke Light Microscopy core facility; and for phalloidin/paxillin/DAPI on ERISM substrates a Nikon Eclipse Ti2 epi fluorescence microscope.

### Quantification of immunofluorescence images

Fluorescence intensity was quantified via the ImageJ software. For the intensity of actin and paxillin, a 50 pixel of rolling ball radius background was subtracted. The actin intensity was normalized to the used exposure time (1 - 2 s for siScr and 1 s for si*CRB2*).

The calculation of the actin orientation order parameter was adapted from Ref. (64). The quantification of counts and mean size of paxillin stains was adapted from Ref. (65). In brief, background subtraction was performed as described above. The local contrast of the images was enhanced by the CLAHE (Contrast Limited Adaptive Histogram Equalization) plug-in using (“block size” = 19, “histogram bins” = 256, “maximum slope” s= 6, “no mask and fast”). The background was minimized by the “Exponential” command. After auto-adjusting brightness and contrast, the Mexican Hat Filter plug-in was used with radius = 3.5. Next, a threshold of 0 - 255 was applied on the images with the “Default” method and with selecting “Calculate threshold for each image”. Finally, the “Analyze Particles” command was executed with a “size (μm^2^)” parameter of 0.01 - 15.

### Western blot analysis

Following treatment, mature immortalized murine podocyte cultures were washed once with ice cold PBS. Cells were then harvested in loading buffer (Cell Signaling Technologies; Boston, MA, USA) supplemented with 1 µM Calyculin A (EMD Chemicals; Billerica, MA, USA) and protease inhibitor cocktail, 1:400 dilution (Sigma). Whole cell extracts were passed through a 1cc syringe (BD Biosciences; San Jose, CA, USA) ten times. Cell lysates were then subjected to SDS-polyacrylamide gel electrophoresis using NuPAGE 10% Bis-Tris pre-cast gels (Invitrogen), followed by transfer to 0.2 µM pore size PVDF membrane (Millipore; Billerica, MA, USA). Membranes were blocked with 1 hour incubation in 5% milk in PBS. Protein immunoblotting was performed using phospho-paxillin (Y118) rabbit polyclonal antibody (1:1000, Cell Signaling Technologies, 2541), phospho-FAK (Y397) rabbit monoclonal (D20B1) antibody (1:1000, Cell Signaling Technologies, 8556) and total FAK rabbit monoclonal antibody (1:1000, Cell Signaling Technologies, 71433). Membranes were washed once with PBS and incubated for 1 hour with horse radish peroxidase-conjugated goat anti-rabbit polyclonal secondary antibody at 1:10,000 dilution (Invitrogen). Immuno-labeled proteins were detected using a chemiluminescence detection system (Pierce Biotechnology; Rockford, IL, USA) on Kodak Biomax film (VWR Scientific; Radnor, PA, USA).

### Luciferase reporter assay

Cells were plated in triplicate at 2 x 10^5^ cells/ml in 2 ml on 6-well collagen coated plates. The plates were incubated at 33 °C and 5% CO_2_ until 95% confluency was reached. Protein lysates were obtained by removing the media and rinsing cells twice with 1 ml 1xPBS. Cells were then incubated in 200 µl of 1x Passive Lysis Buffer (PLB, Promega E1500) on a 150 RPM orbital shaker for 45 min. Lysates were collected in a microcentrifuge tube and centrifuged at 800 RPM for 5 min at 4 °C. The supernatant was transferred to a clean microcentrifuge tube and stored on ice. Quantification of the whole protein lysates was performed using the BioRad Protein Assay (cat. #5000001) using 0.5 mg/ml to 0.05 mg/ml of BSA standards and the Microtiter Plate protocol. Each standard sample was run in triplicate with 10 µl of each pipetted into 200 µl of diluted 1x Dye Reagent. The plate was incubated for 50 min at room temperature and then the absorbance at 595 nm was read using a microplate reader (M200Pro, Tecan). The absorbance of the samples was plotted against the standard curve with an *R*^2^ value of > 0.98 to measure the concentration of each protein sample. Each sample was diluted to 0.35 mg/ml to perform the Luciferase Assay (Promega E1500). Luciferase were thawed in a room temperature water bath, and the Luciferase Assay Buffer was added to the lyophilized Luciferase Assay Substrate. Each sample was plated in triplicate, along with No Template Controls containing only 1x PLB in a 3990 Costar Plate. 20 µl of normalized protein lysate was added to the plate, and 100 µl of diluted Luciferase Assay Reagent were added immediately before loading samples into the microplate reader to measure luminescence (integration time, 10 s).

### ELISA assay

Immunoassays were used to quantify concentrations of connective tissue growth factor (CTGF) (Abcam, ab261851), Thrombospondin-1, Endothelin-1, Cyr61/CCN1 (R&D Systems; DTSP10, DET100, DCYR10, respectively), and transforming growth factor-beta 1 and 2 (TGF-β1, -2, and -3) (MesoScale Discovery, K15241K), in cell culture supernatants. Due to all samples having concentrations below the lowest limit of detection for the assay, TGF-β3 was excluded from the analysis. CTGF required a 10-fold dilution of supernatant and had a mean intra-assay CV of 1.6%. Thrombospondin (diluted 2-fold), Endothelin-1 (diluted 10-fold), and Cyr61 (undiluted) had mean intra-assay CVs of 1.3%, 2.1%, and 3.1%, respectively. All samples required acid treatment for the TGF-β assay. Ten microliters of 1 M HCl was added to 50 µl of sample and incubated for 10 minutes after which 7 µl of 1.2 M NaOH was added before vortexing the sample and loading onto the plate. All assays were performed according to manufacturer’s instructions.

### Elastic resonator interference stress microscopy

ERISM substrates were fabricated as described earlier (43). In brief, a bottom mirror, consisting of a 0.5-nm-thick Cr adhesive layer, a 10-nm-thick Au mirror layer and a 50-nm-thick Ta_2_O_5_ protective layer, were deposited on clean glass coverslips (No.5) by electron beam evaporation, thermal evaporation and sputtering, respectively. The siloxane-based elastomer (GEL8100, Nusil) was prepared in a 1:1 mass ratio and spin-coated onto the bottom mirror. The top surface of the elastomer was treated with 2.5 – 15% power of oxygen plasma (max. power, 600 W; duration, 10 s) to reduce its surface energy and improve the quality of the 15-nm-thick Au mirror deposited on top.

A set of 4 silicon chambers (surface area each, 0.75 × 0.75 cm^2^; 81201, Ibidi) were applied on the substrate. The chip surface was coated with 8 μg/cm^2^ of type I collagen (Rat Tail, A10483-01, Gibco) for one hour at room temperature. Chambers were washed three times with 1xPBS and three times with fresh cell medium (RPMI 1640/ 10% FBS/ 1% PS/ 1% ITS), and then incubated at 33 °C, 5% CO_2_ for 30 min. Trypsinized podocytes were seeded at a density of 1000 cells per chamber. Podocytes were incubated at 33 °C, 5% CO_2_ for 24 h before transfer to 37 °C, 5% CO_2_ for the 12-day differentiation. For the maintenance of cell culture on ERISM substrates, the cell medium was exchanged daily by washing each chamber three times with pre-warmed fresh cell medium.

For ERISM measurements, the podocyte cultures were placed in a microscope on-stage incubator (Okolab) and maintained at 37 °C, 5% CO_2_. The incubator was mounted on a modified inverted fluorescence microscope (Nikon Ti2). Monochromatic probe light with wavelengths ranging from 550 to 750 nm was provided by a halogen lamp (ASBN Series, Oceanhood) filtered by a ⅛ m monochromator (CM110, Spectral Products). The probe light was coupled to the microscope and projected onto the substrate with either a 20×/0.45 NA or 40×/0.60 NA objective (Nikon S Plan Fluor) depending on the required field of view and magnification. Reflection and phase contrast images of the sample were detected by an sCMOS camera (0.05 s exposure time; Zyla, Andor Technology) and a standard CMOS camera (iCube C-NS4133BU, NET GmbH), respectively. The local resonance wavelengths of the sensor for each cell were measured by recording wide-field reflectance images for a series of illumination wavelengths. The wavelength selection and image capturing were controlled by an in-house developed software (LabView, NI) written by Phillip Liehm.

Reflectance images were converted into local displacement maps by comparing the local resonance wavelengths against the predictions of an optical model of the ERISM substrate for different elastomer thicknesses as described previously (43, 63). The Fourier-filtered ERISM displacement maps were obtained by targeting spatial frequencies in the range of 0.016 – 1.007 µm⁻¹ in the obtained displacement maps using the FFT plugin of the ImageJ software.

### Statistical analysis

Data groups for both cell lines were compared using two sample or paired-sample student *t*-test (equal variance not assumed). *P*-values of statistical significance are represented as: n.s.: *p* > 0.05, ∗: *p* ≤ 0.05, ∗∗: *p* ≤ 0.01, ∗∗∗: *p* ≤ 0.001.

## RESULTS

### Family Clinical Characteristics

Family DUK40595 is a 2-generation East Asian kindred from India with two affected offspring from a non-consanguineous marriage (Figure 1a). A summary of all relevant clinical information for the family is provided in Table 1. The proband (DUK40595 – 1) and her affected male sibling (DUK40595 – 100) developed SRNS in their first decade of life (Table 1). The parents are healthy with no evidence of kidney disease. The female sibling was the only member of the family to undergo kidney biopsy which was diagnostic for SRNS/FSGS (Table 1). Subsequent clinical genetic testing of the proband revealed a compound heterozygous mutation of *CRB2* comprised of the previously reported 16-bp duplication p.Gly1036_Alafs*43 mutation (23) and a rare in-frame, 9-bp deletion mutation (c.3220_3228del (p.Leu1074_Asp1076del); minor allele frequency - 0.00004). The duplication results in a frameshift and premature truncation of the protein 43 amino acids downstream of codon 1036 within the third Laminin G domain (p.Gly1036Ala*43; ENST00000373631) (Figure 1b). In addition, a second rare, heterozygous 9-bp deletion in exon 10 of *CRB2* (chr9:126136027_126136035del; Depth - 167x) was identified. The 9-bp deletion results in the loss of three highly conserved amino acid residues in the 12^th^ EGF-binding domain (Figure 1b). The Asp-Leu-Phe residues are conserved to zebrafish (Figure 1c) and their deletion is predicted to disrupt the secondary structure of the 12^th^ EGF-binding domain (Figures 1b and d).

**Figure 1:**
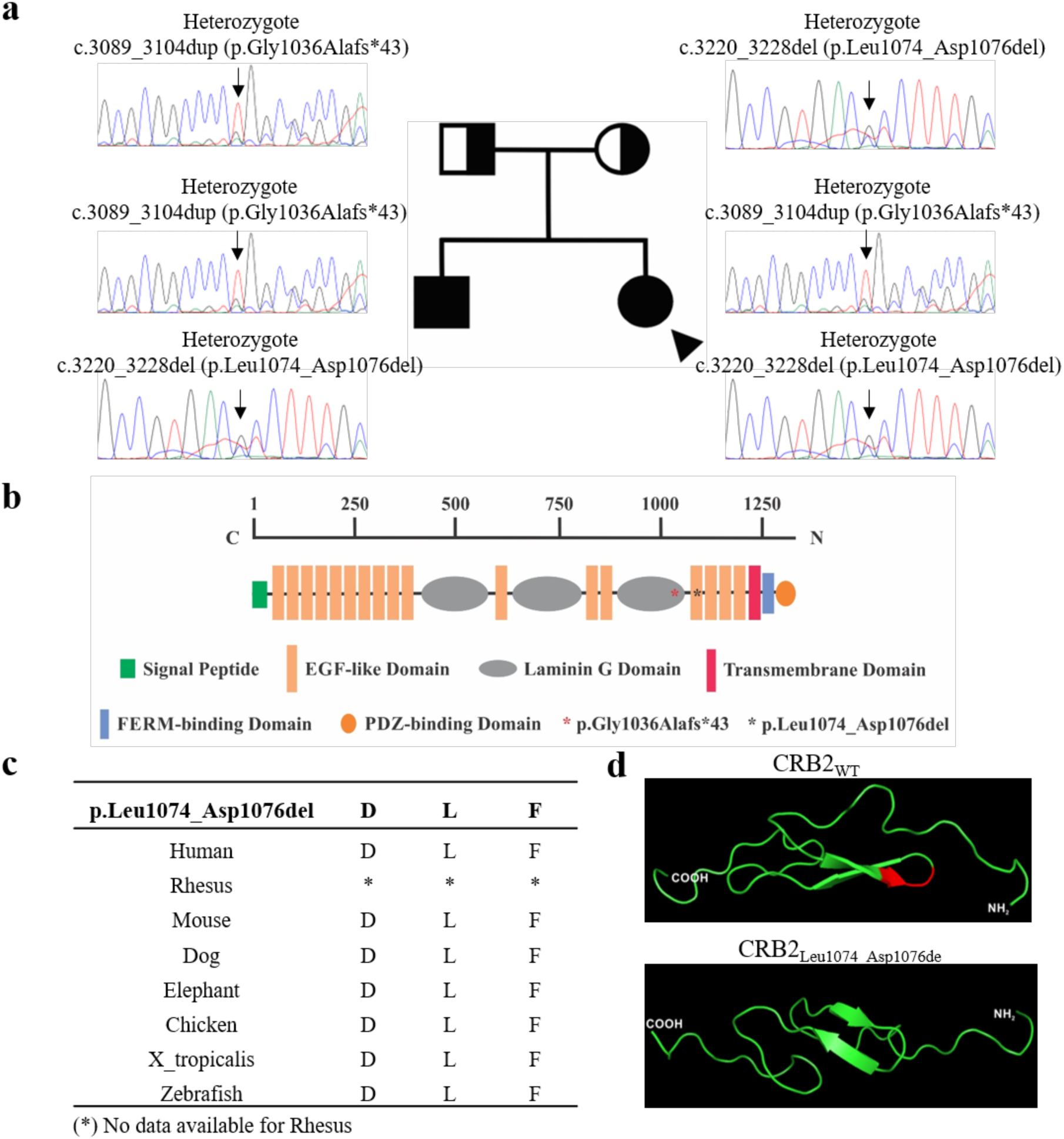
Pedigree of family DUK40595 and identification of CRB2 variant. a) Family DUK40595 is a 2-generation Asian kindred from India. The proband (arrow head) has biopsy-proven FSGS and her younger male sibling has proteinuric nephropathy likely caused by the FSGS lesion. The cause of FSGS in both affected siblings was found to be a compound heterozygous mutation of *CRB2* (FSGS-9). The pathogenic mutation is comprised of a previously described truncating frameshift mutation (p.Gly1036_Alafs*43) and a rare 9-bp deletion mutation (p.Leu1074_Asp1076del). b) Schematic of CRB2 protein functional domains. Family DUK40595 pathogenic mutations identified in the 3rd Laminin G domain (p.Gly1036Alafs*43, [* in orange]) and the 12th EGF-like domain (p.Leu1074_Asp1076del, [* in black]). c) The 9-bp, in-frame deletion results in the loss of three highly conserved amino acids (Asp-Leu-Phe) d) which significantly distorts the structure of the EGF-like domain.

**Table 1:**
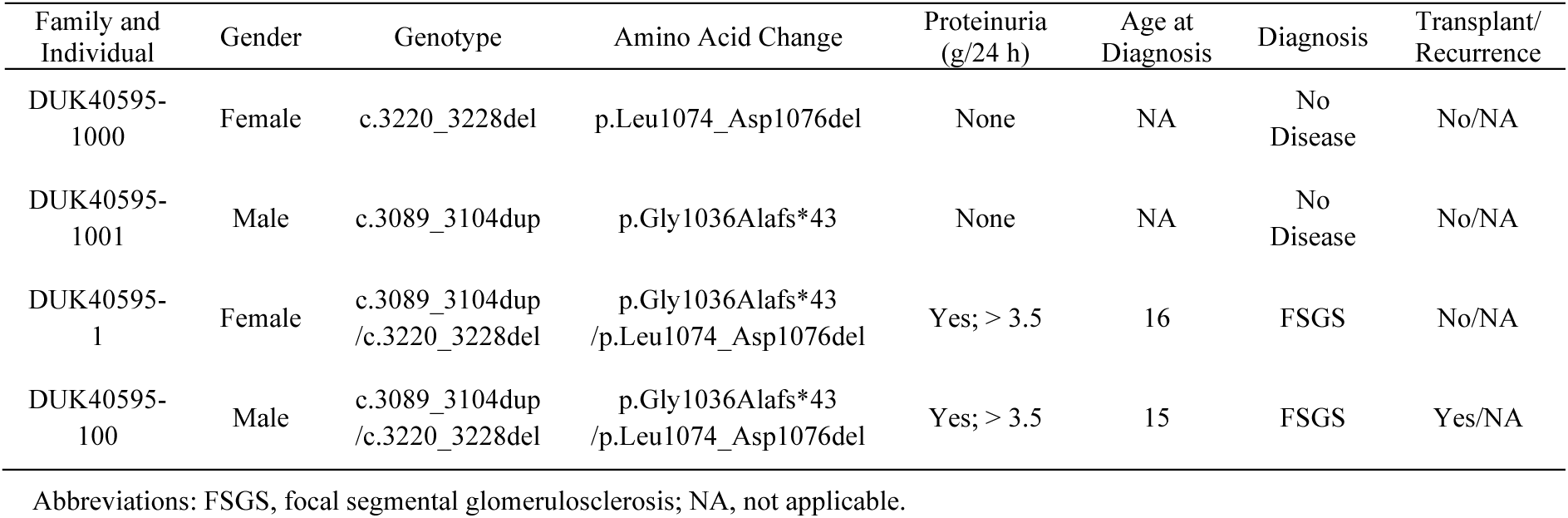
Family DUK40595 Clinical Characteristics

| Family and Individual | Gender | Genotype | Amino Acid Change | Proteinuria (g/24 h) | Age at Diagnosis | Diagnosis | Transplant/Recurrence |
| --- | --- | --- | --- | --- | --- | --- | --- |
| DUK40595-1000 | Female | c.3220_3228del | p.Leu1074_Asp1076del | None | NA | No Disease | No/NA |
| DUK40595-1001 | Male | c.3089_3104dup | p.Gly1036Alafs*43 | None | NA | No Disease | No/NA |
| DUK40595-1 | Female | c.3089_3104dup<br>/c.3220_3228del | p.Gly1036Alafs*43<br>/p.Leu1074_Asp1076del | Yes; > 3.5 | 16 | FSGS | No/NA |
| DUK40595-100 | Male | c.3089_3104dup<br>/c.3220_3228del | p.Gly1036Alafs*43<br>/p.Leu1074_Asp1076del | Yes; > 3.5 | 15 | FSGS | Yes/NA |

Abbreviations: FSGS, focal segmental glomerulosclerosis; NA, not applicable.

Direct sequencing of all family members confirmed the compound heterozygous *CRB2* mutation in the proband and her male sibling. The father (DUK40595 – 1001) was found to be heterozygous for the p.Gly1036-Alafs*43 mutation and the mother (DUK40595 – 1000) was heterozygous for the p.Leu1074_Asp1076del mutation (Table 1 and Figure 1). The male sibling eventually progressed to ESKD at age 17 and received a kidney transplant without recurrence of disease in his allograft.

### *CRB2* Knockdown Alters Podocyte Spreading Area and Maturity Marker Expression

Prior studies of the p.Gly1036Alafs*43 mutation have shown that it functionally eliminates the CRB2 transmembrane domain and markedly reduces glomerular CRB2 expression (Figure S2) (23). To evaluate the effects of CRB2-deficiency on podocyte mechanosignaling, we generated conditionally immortalized human podocytes stably-expressing *CRB2* siRNA (si*CRB2*) and compared them to controls expressing a scrambled RNA segment of equivalent length (siScr) (66). Both cell lines contained human telomerase reverse transcriptase (hTERT) and a temperature-sensitive SV40 transgene. The podocytes were initially cultured under growth-permissive conditions and then differentiated over the course of 12 days under growth restrictive conditions. In differentiated podocytes, we achieved an ∼80% reduction in *CRB2* mRNA expression in the si*CRB2* line relative to controls (siScr), as shown in Figure 2a.

**Figure 2:**
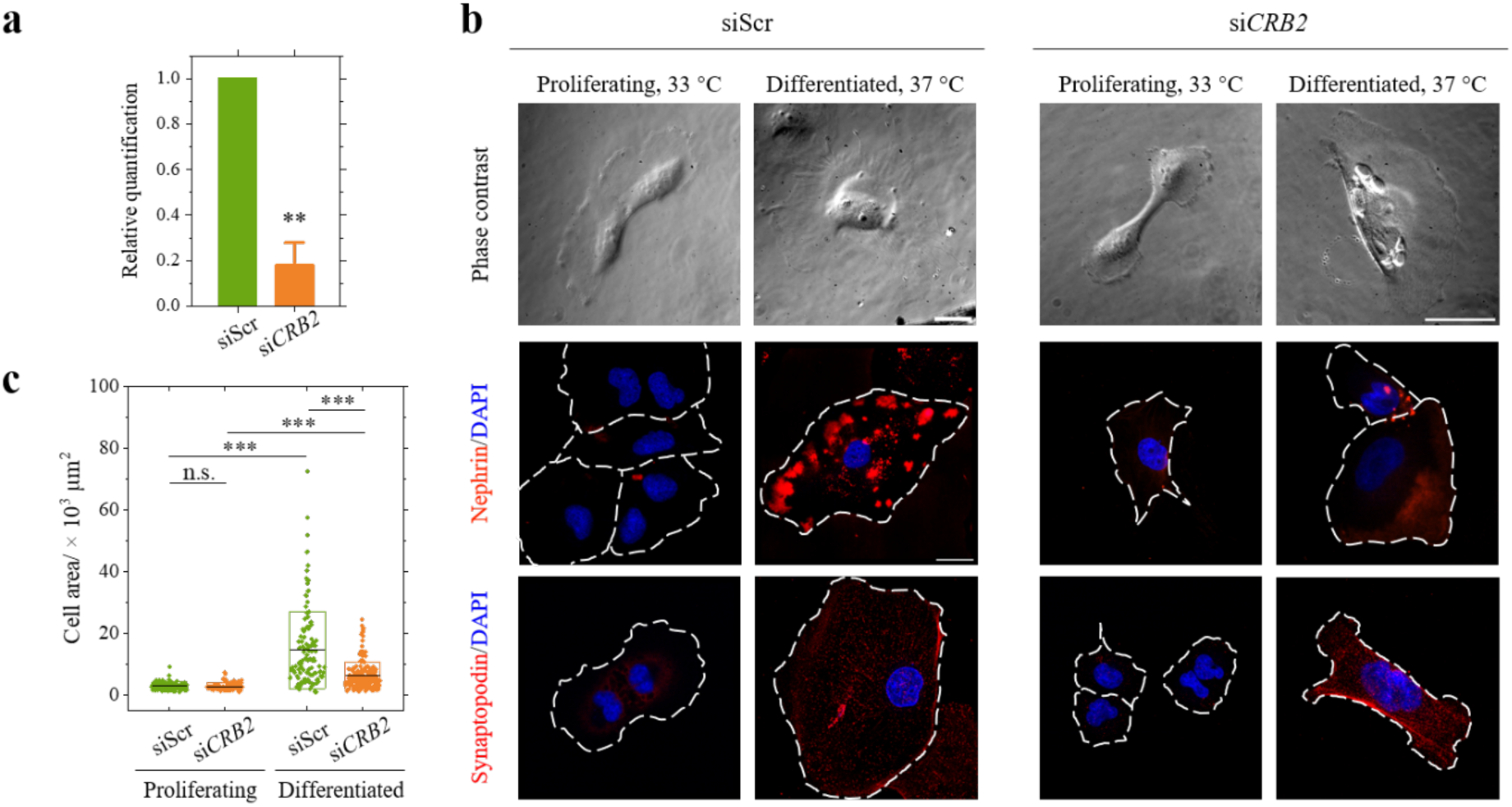
Differentiation of human immortalized urinary podocytes. **a)** Plot of relative quantification of *CRB2* gene expression in scrambled siRNA (siScr) and *CRB2*-knockdown (si*CRB2*) podocytes from three quantitative polymer chain reaction (qPCR) replicates. Mean values (boxes) and standard deviation (error bar). Groups were compared using a paired-sample *t*-test with non-equal variance, **: *p* ≤ 0.005. **b)** Phase contrast and immunofluorescence images of nephrin (red), synaptopodin (red) and nuclei (detected by DAPI, blue) for proliferating (at 33 °C) and 12-day-differentiaed (at 37 °C) siScr and si*CRB2* podocytes. Cell outlines are indicated by white-dashed lines in the immunofluorescence images. Scale bars, 50 μm. **c)** Comparison of cell area between proliferating and 12-day-differentiated podocytes (dots), showing mean (lines) and ±1 SD (boxes) for the siScr and si*CRB2* cells. Groups were compared using two-tailed *t*-tests with non-equal variance, n.s.: *p* > 0.05, *: *p* ≤ 0.05, **: *p* ≤ 0.01, ***: *p* ≤ 0.001.

Figure 2b shows phase contrast images of siScr and si*CRB2* podocytes under growth-permissive and growth-restrictive conditions. The quantification of cell area in Figure 2c indicates that both cell lines increased their spreading area during differentiation. However, the spreading area was roughly half as much for the *CRB2* knockdown podocytes compared to the siScr cells (siScr: mean cell spreading area: proliferating - 3,032 µm^2^, differentiated - 14,638 µm^2^; si*CRB2*: mean cell area: proliferating - 2,862 µm^2^, differentiated - 6,397 µm^2^).

We further assessed differentiated podocytes for expression and localization of the slit diaphragm protein nephrin and the foot process protein synaptopodin looking at both proliferating and differentiated podocytes (Figure 2b). Confocal imaging of podocytes revealed a marked increase in both nephrin and synaptopodin expression in differentiated cells for both lines. However, nephrin staining appeared to be reduced in si*CRB2* podocytes relative to controls, consistent with prior observations (26, 32).

Taken together, CRB2 knockdown reduces the podocyte spreading area in differentiated cells and alters the expression of nephrin.

### *CRB2* Knockdown Induces YAP Target Gene and Protein Expression in Immortalized Podocytes

In order to investigate if *CRB2* knockdown alters YAP activity in differentiated podocytes, we performed Transcriptional Enhance Associate Domains (TEAD) luciferase reporter assays using the well-validated 8xGTIIC-luciferase reporter (67). We detected high YAP activity in both lines; however, TEAD reporter activation was significantly increased in si*CRB2* podocytes relative to control (Figure 3a). Additionally, we performed Western Blot analyses to evaluate YAP phosphorylation at serine 127 (Ser127). Phosphorylation of YAP at Ser127 promotes its degradation or cytoplasmic sequestration through binding with 14-3-3 family proteins. Cytosolic retention of the co-transcriptional regulator renders it inactive and prevents target gene activation (68). Conversely, when YAP is not phosphorylated at serine 127, it correlates with increased nuclear translocation and transcriptional activity. In Figures 3b and 3c, we show that transcriptionally active non-phosphorylated YAP (np-YAP) is significantly upregulated in the si*CRB2* line relative to controls, which correlates with a marked enhancement of thrombospondin-1 (THBS-1, Figure 3d), a YAP target gene known to drive mechanosignaling via direct stimulation of the integrin receptor (69, 70). Finally, to more broadly assess the effects of CRB2 deficiency on YAP mechanosignaling target gene expression, we performed ELISA assays and demonstrate significantly increased release of the YAP target genes Transforming Growth Factor-β1 (TGF-β1), Transforming Growth Factor-β2 (TGF-β2), THBS-1, Connective Tissue Growth Factor/CCN Family Member 2 (CTGF/CCN2), Cysteine-rich Angiogenic Inducer 61/CCN Family Member 1 (Cyr61/CCN1) and Endothelin-1 (EN1) from si*CRB2* podocytes relative to controls (Figure 3e).

**Figure 3:**
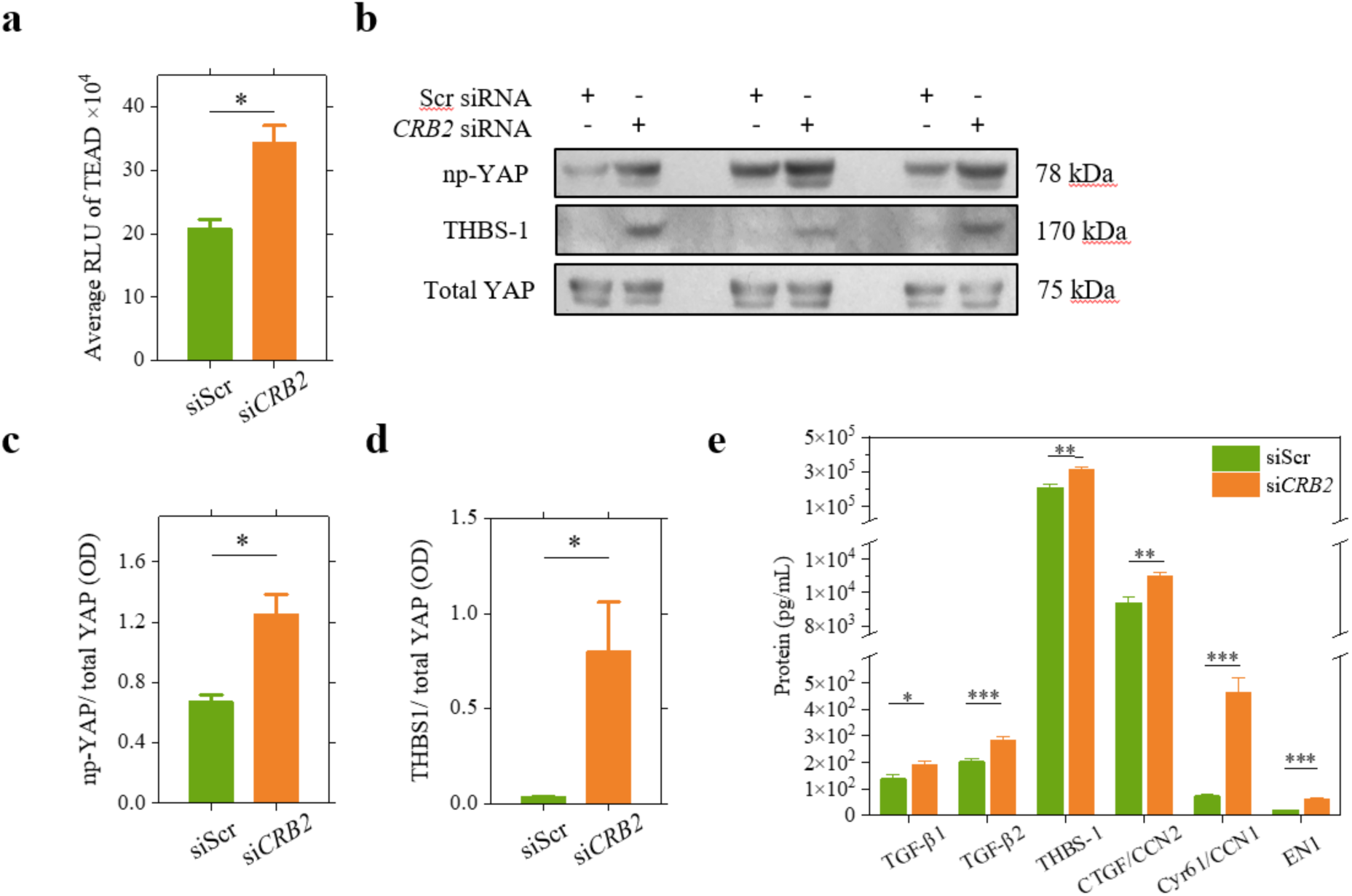
Upregulated YAP signaling in human podocytes upon *CRB2* knockdown. **a**) Luciferase reporter assay of average expression of transcriptional enhanced associate domain (TEAD) in siScr and si*CRB2* podocytes. *N* = 6. **b**) Western blots of non-phosphorylated YAP (np-YAP), Thrombospondin-1 (THBS-1) and total YAP expression in siScr and si*CRB2* podocytes. **c**) and **d**) Relative expression level of np-YAP and THBS-1 compared to total YAP for both cell lines using data obtained from b). **e**) Enzyme-linked immunoassay (ELISA) analysis of expression of YAP-target genes for both cell lines. TGF, transforming growth factor; CTGF, Connective tissue growth factor; Cyr61, Cysteine-rich angiogenic inducer 61; CCN1/2, Cellular communication network factor 1 or 2; EN1, Endothelin 1. *N* = 6. Mean values (boxes) and ±S.E.M. (error bars). Groups in panels a), c), d), and e) were compared using two-tailed *t*-tests with non-equal variance, n.s.: *p* > 0.05, *: *p* ≤ 0.05, **: *p* ≤ 0.01, ***: *p* ≤ 0.001.

Taken together, these data suggest that *CRB2* knockdown induces YAP dephosphorylation/activation and mechanotransduction gene expression in podocytes.

### *CRB2* Knockdown Enhances Podocyte Contractile Force Development and Contractility ***in vitro***

YAP is a key regulator of cellular mechanotransduction in multiple cell types (67, 71). Prior studies have shown that the YAP target genes *CTGF/CCN2*, *Cyr61/CCN1* and *THBS-1* are all direct activators of mechanosignaling through integrin receptors (72–75). Downstream effectors of integrin receptor signaling, such as focal adhesion kinase (FAK) and the adaptor protein paxillin, populate focal adhesions and transmit outside-to-inside signals to direct various aspects of cellular motility and attachment (76, 77). Since CRB2 knockdown induces YAP activity and the expression of YAP mechanotransduction target genes, we next explored the effect of CRB2 knockdown on podocyte contractile force development using ERISM.

ERISM detects cell-induced deformations of an elastic, optical micro-cavity by detecting and analyzing the interference of monochromatic probe light across a field of view captured by a modified inverted microscope (43, 47). We have previously utilized ERISM to observe the near-complete loss of immortalized human podocyte forces in a puromycin amino nucleoside (PAN) injury model (78).

To characterize the effects of CRB2 knockdown on podocyte force development, isolated siScr and si*CRB2* podocytes were seeded on collagen-I-coated ERISM substrates with an apparent stiffness ranging from 11 – 142 kPa, and ERISM measurements for *n* ≥ 14 cells were performed on 4 or 5 non-consecutive days during the 12 days of podocyte differentiation. Figures 4a and 4b show representative phase-contrast and ERISM displacement images of both cell lines when using a substrate with an apparent mechanical stiffness of 74 kPa. Contractile forces generated inside the cell result in an expansion of the elastomer at the position of focal adhesions (FA; indicated by bright areas at the cell periphery) (44). At the same time, cell contraction also leads to a compression of the elastomer underneath the cell body (indicated by the dark regions). These vertical deformation patterns are a response of the deformable substrate to predominantly horizontally directed cellular forces (43). (For a more-detailed discussion of force transmission in podocytes, see Figure 6 below and Ref. (78).)

**Figure 4:**
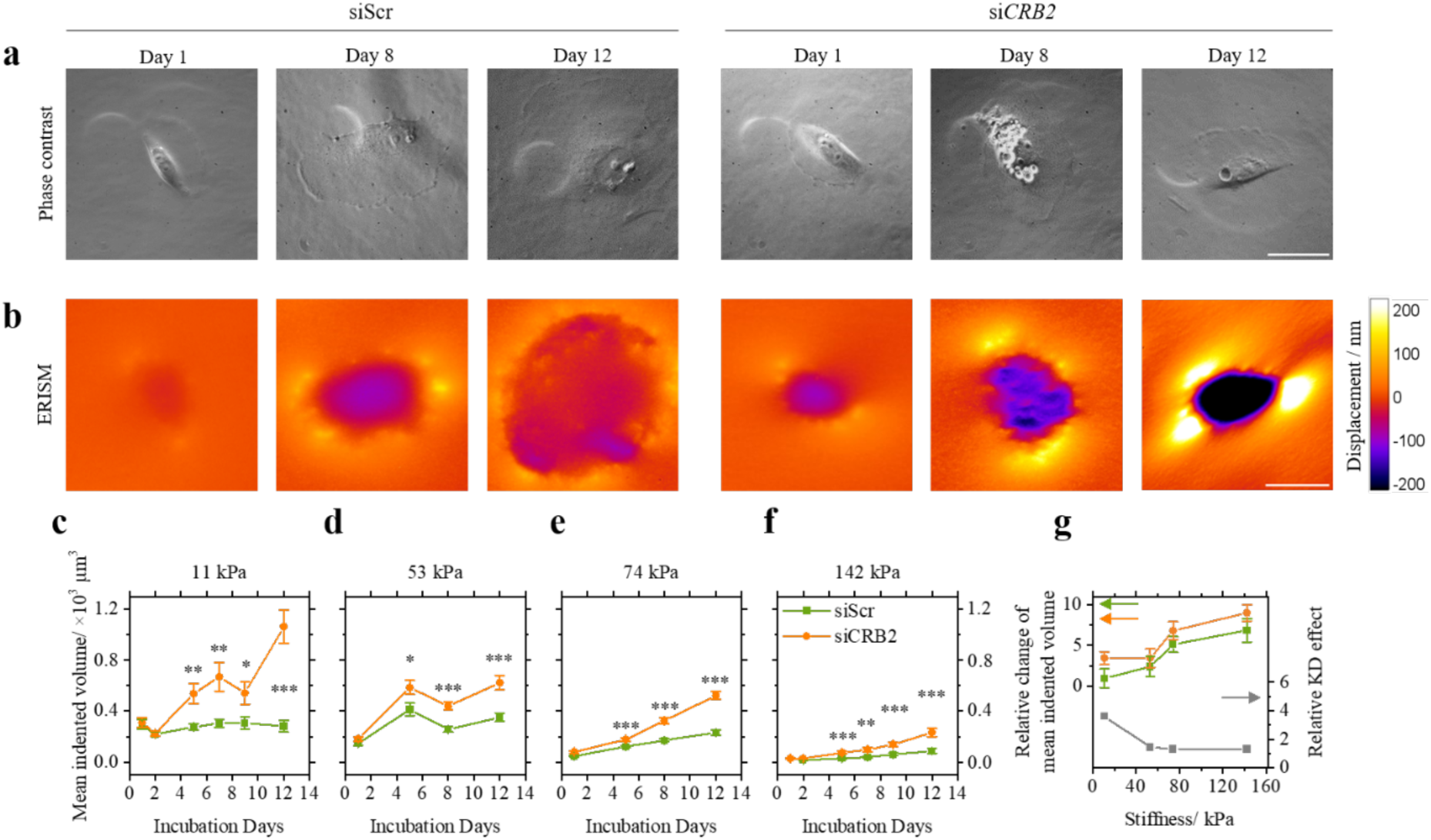
Cell contractility of si*CRB2* podocytes increases throughout differentiation. **a)** Phase-contrast images and **b)** ERISM displacement maps of representative control (siScr) and *CRB2* knock-down (si*CRB2*) podocytes on ERISM substrates with an apparent stiffness of 74 kPa on Day 1, 8 and 12 of the differentiation at 37 °C. Scale bars, 50 μm. **c-f)** Evolution of mean indented sensor volume (symbols) and ±S.E.M. (error bars) of siScr (green) and si*CRB2* (orange) podocytes during differentiation, measured on ERISM substrates with an apparent stiffness of 11, 53, 74 and 142 kPa, respectively. *N* ≥ 20 for each group. **g)** Relative change of mean indented volume between Day 12 and Day 1 of differentiation (symbols) and ±SD (error bars) of siScr (green) and si*CRB2* (orange) podocytes. Relative knockdown (KD) effect, i.e. the ratio of relative change, si*CRB2*/siScr cell line, for ERISM substrate of different stiffness (grey symbols, right y-axis). Groups were compared using two-tailed *t*-tests with non-equal variance, *: *p* ≤ 0.05, **: *p* ≤ 0.01, ***: *p* ≤ 0.001.

The magnitude of the exerted contractile podocyte forces was quantified by calculating the total volume of downward sensor deformation underneath the cell soma, referred to as the indented volume (*IV*) in the following. Figure 4c-f show the *IV* for si*CRB2* and siScr podocytes during the differentiation period when cultured on substrates with different mechanical stiffnesses and demonstrate that cell contractility generally increases during differentiation. From Day 4 of the differentiation onwards, the contractility of si*CRB2* podocytes proved significantly higher than that of siScr controls, regardless of substrate stiffness. These findings suggest that *CRB2* knockdown significantly enhances podocyte contractility during differentiation.

Next, we calculated the relative change in *IV* during differentiation, i.e., between Day 12 and Day 1 or 2 of the differentiation, for all tested substrates (Figure 4g), and found that it increases with substrate stiffness for both cell lines. In order to evaluate, if the mechanosensing capability of CRB2 knockdown cells is changed, we normalized the relative change in *IV* during differentiation of si*CRB2* podocytes to the value of siScr controls (‘relative knockdown effect’ in Figure 4g). Strikingly, the relative effect of CRB2 knockdown is largest for the softest substrate and decreases with increasing substrate stiffness. This indicates that the contractility of CRB2 knockdown cells remains high on soft substrates compared to control cells, which implies that mechanosensing might be impaired in CRB2 knockdown cells, rendering them unable to reduce contractility on highly compliant substrates. This finding is in line with the suggested role of CRB2 as an upstream regulator of YAP activity.

Collectively, our data from this series of experiments show that CRB2 knockdown increases force generation in human podocytes. Since this effect is the highest on soft substrates, CRB2 may function as an upstream regulator of YAP-induced mechanoactivation.

### CRB2 Knockdown Enhances Filamentous Actin and Mechanosensitive Protein Expression in Podocytes

Because CRB2 knockdown increases podocyte force development, we wanted to determine whether this enhancement in force is associated with an increase in the expression of mechanosensitive proteins. To evaluate this, we performed immunocytochemistry in si*CRB2* and siScr podocytes fixed on ERISM substrates with an apparent stiffness of 74 kPa. First, we explored the expression and localization of F-actin and the FA adaptor protein paxillin (Figure 5a). While actin stress fibers are visible in both cell lines, quantifying the mean fluorescent intensity of F-actin shows a significant increase of F-actin in differentiated si*CRB2* podocytes as compared to the siScr control group (Figure 5c). This finding is in contrast to results from an earlier study that found a reduction in F-actin formation for *CRB2* knockout podocytes (27). Additionally, to test if the CRB2 knockdown leads to a rearrangement of the actin cytoskeleton, we calculated the orientation order parameter *S* of actin fibers (Figure 5a; *S* = 0 for isotropic, non-polarized; *S* = 1 for perfectly linearly polarized actin cytoskeleton) (64). The orientation order of si*CRB2* podocytes displayed a slight increase compared to siScr controls, yet with no statistical significance (Figure 5d).

**Figure 5.**
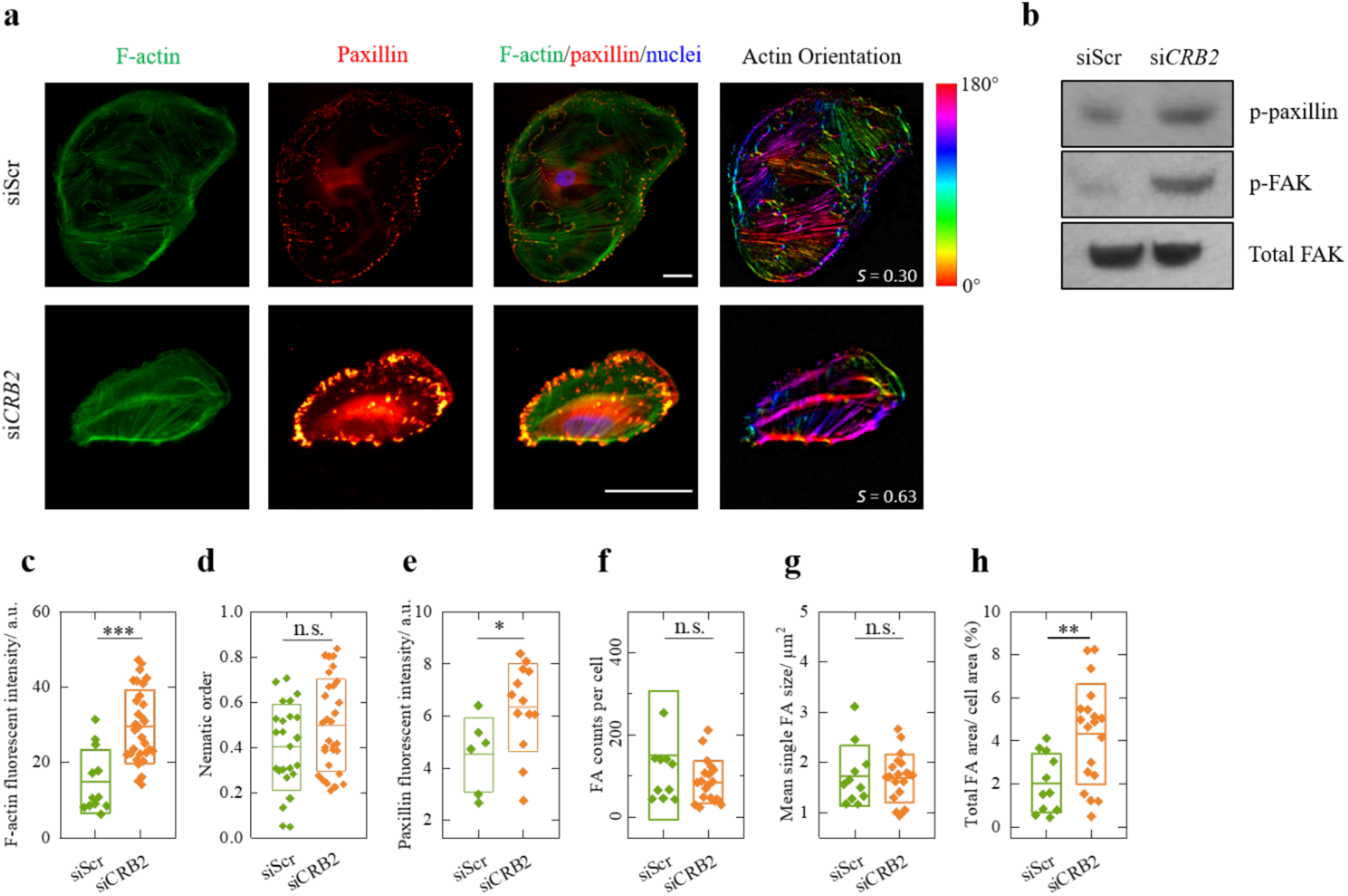
Mechano-sensitive proteins in differentiated human podocytes. **a)** Representative immunofluorescence images of F-actin (labelled by TRITC-phalloidin, green), paxillin (red) and nuclei (labelled by DAPI, blue) for 12-day-differentiated control (siScr) and *CRB2* knockdown (si*CRB2*) podocytes. Scale bars, 50 μm. F-actin, filamentous actin; YAP, yes-associated protein. **b)** Western blot of phosphorylated paxillin, focal adhesion kinase (p-paxillin, p-FAK) and total FAK in siScr and si*CRB2* podocytes. Since the experiment was not repeated, we did not perform quantitative analysis. Mean fluorescence intensity of **c)** F-actin and **e)** paxillin per cell for both cell lines. **d)** Actin cytoskeleton order parameter (nematic order, *S*) for both cell lines. *S* = 0 for isotropic, non-polarized and *S* = 1 for perfectly linearly polarized actin cytoskeleton. **f)** Numbers of focal adhesion (FA) clusters per cell (quantified via the counts of paxillin dots in immunofluorescence images), **g)** mean size of individual FAs and **h)** ratio of total FA area across a cell to its size for differentiated siScr and si*CRB2* podocytes. Shown are the values for individual cells (symbols), the mean (central lines) and ±SD (boxes). At least 6 cells were measured per group. Groups were compared using two-tailed student *t*-tests with non-equal variance, n.s.: no significance, *: *p* ≤ 0.05, **: *p* ≤ 0.01, ***: *p* ≤ 0.001.

In both cell lines, paxillin showed a pronounced dotted/punctuate expression pattern along the cell periphery (Figure 5a), consistent with its position within the FAs of cells (79, 80). No difference in the distribution of paxillin was observed between the cell lines. However, western blotting of phosphorylated paxillin (p-paxillin) and focal adhesion kinase (p-FAK, Figure 5b), as well as the immunolabelling of FAK (Figure S3) suggest the presence of enhanced paxillin and FAK activity in CRB2 deficient cells. Assuming that each paxillin dot in the immunofluorescence images corresponds to a FA, we quantified the number of FAs per cell; on average 150 and 85 FAs were found in siScr and si*CRB2* podocytes, respectively (Figure 5f). The average size of individual FAs, approximately 1.7 µm² (Figure 5g), was independent of cell type and consistent with earlier measurements of FA size (80, 81). However, due to the smaller cell area of knockdown cells, the ratio of total FA area to cell area was significantly higher in si*CRB2* podocytes relative to siScr controls (Figure 5h). Since FAs grow in response to increased cell contractility (82), these data suggest that CRB2 knockdown enhances formation of F-actin fibers, FAK and paxillin activation, FA density and contractility in podocytes, probably downstream of YAP target gene expression.

Figure 5a shows that paxillin localizes at the terminus of actin fibers in both cell lines. In order to investigate how contractile forces are transmitted to the substrate at the FA sites in podocytes, we employed Fourier filtering of the ERISM displacement maps. This procedure filters out any broad substrate deformations and thus provides a more detailed view of subtle and localized deformations (Figure 6c). The Fourier-filtered displacement maps show a series of clearly distinguishable push/pull features (shown in red/blue) at the periphery of the cell. This twisting behavior is consistent with earlier observations of the torque being applied by FAs (43). Areas of high paxillin expression are located at the center of these features, confirming that they are associated with FAs transferring traction forces horizontally to the substrate. This twisting is more pronounced in the si*CRB2* cell than the siScr cell, which is in line with the higher overall contractility of the latter (Figure 4) (44, 83).

**Figure 6.**
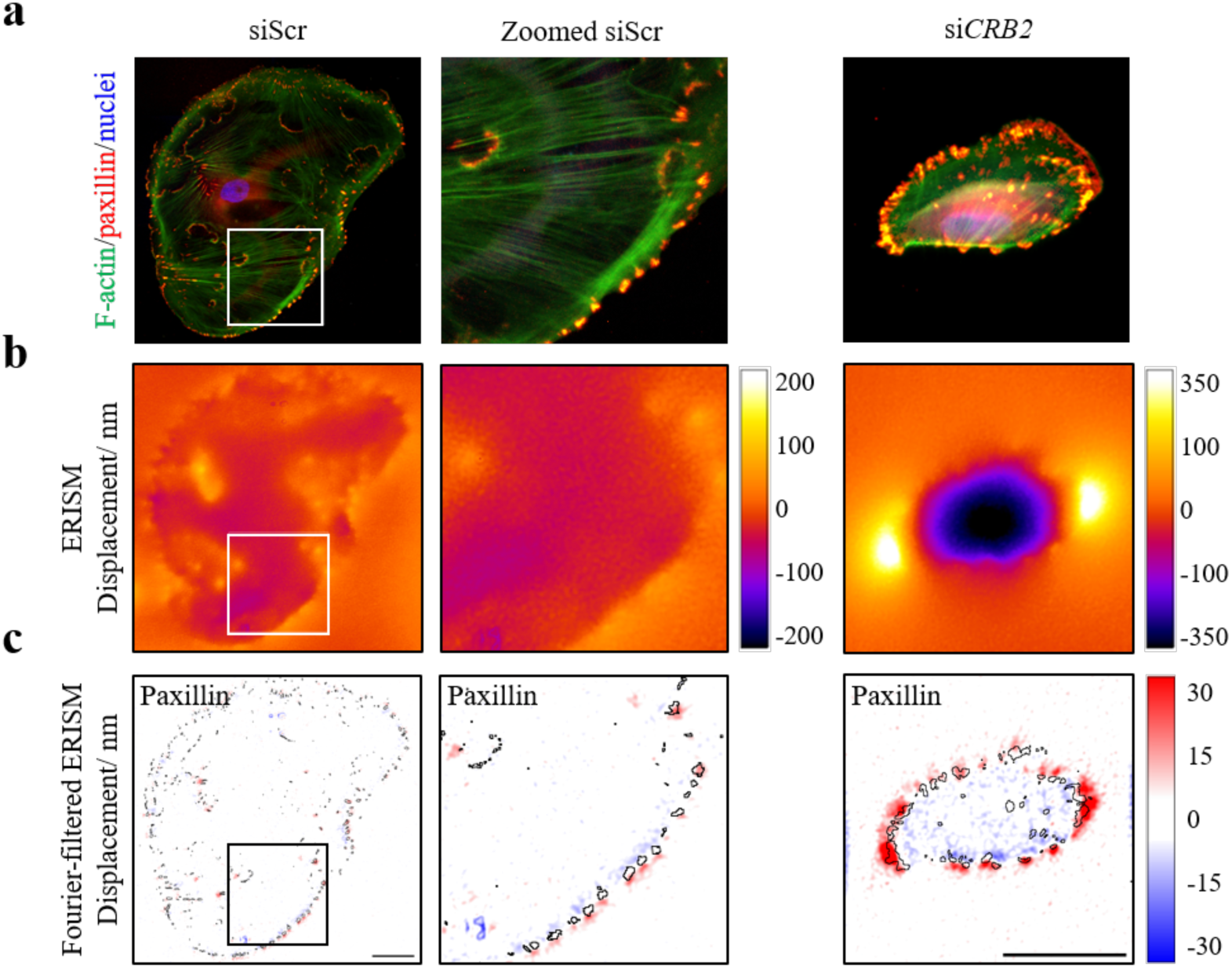
Podocyte contractile forces are colocalized with the paxillin expression FAs at the cell periphery. a) Composite immunofluorescence images of the representative cells shown in Figure 5. F-actin (labelled by TRITC-phalloidin, green), paxillin (red) and nuclei (labelled by DAPI, blue) of a differentiated control podocyte (left), a zoom-in on the area marked by the white square (middle), and same staining for a differentiated si*CRB2* podocyte (right). b) Corresponding ERISM displacement and c) Fourier-filtered ERISM maps, with black lines denoting the outlines of paxillin-rich areas. Scale bars, 50 μm. F-actin, filamentous actin.

### Force Development in CRB2 Knockdown Podocytes is not Significantly Altered by YAP inhibitors or Rapamycin

So far, we have established that CRB2 knockdown leads to upregulation of YAP activity and YAP target gene expression, as well as to an increase in cell forces and an impairment of mechanosensing at low substrate stiffnesses. Next, we asked if the observed mechanical phenotype of CRB2 knockdown cells is lost when the transcriptional activity of YAP is inhibited. To this end we treated differentiated podocytes of both the siScr and si*CRB2* lines with either K-975 (100 nM) or verteporfin (72 – 359 nM; equivalent to 50-250 ng/mL). Both reagents inhibit the interaction between YAP and TEAD (84). However, no significant reduction in cellular forces was observed after treating the cells with either of the two drugs for 24 h; there was also no difference in response between the two cell lines (Figure 7b and 7c). Higher concentrations of verteporfin (1.5 and 5 µM, equivalent to 1.0 and 3.5 µg/mL) did not result in a reduction of the contractility of *CBR2* knockdown cells either (Figure S4).

We also investigated the effect of the mTOR inhibitor rapamycin on the contractility of matured si*CRB2* and siScr podocytes as rapamycin might enhance the autophagy-mediated degradation of YAP (85, 86). Again, no effect on the contractility of either cell line was found after 24 h of treatment (Figure 7d).

**Figure 7.**
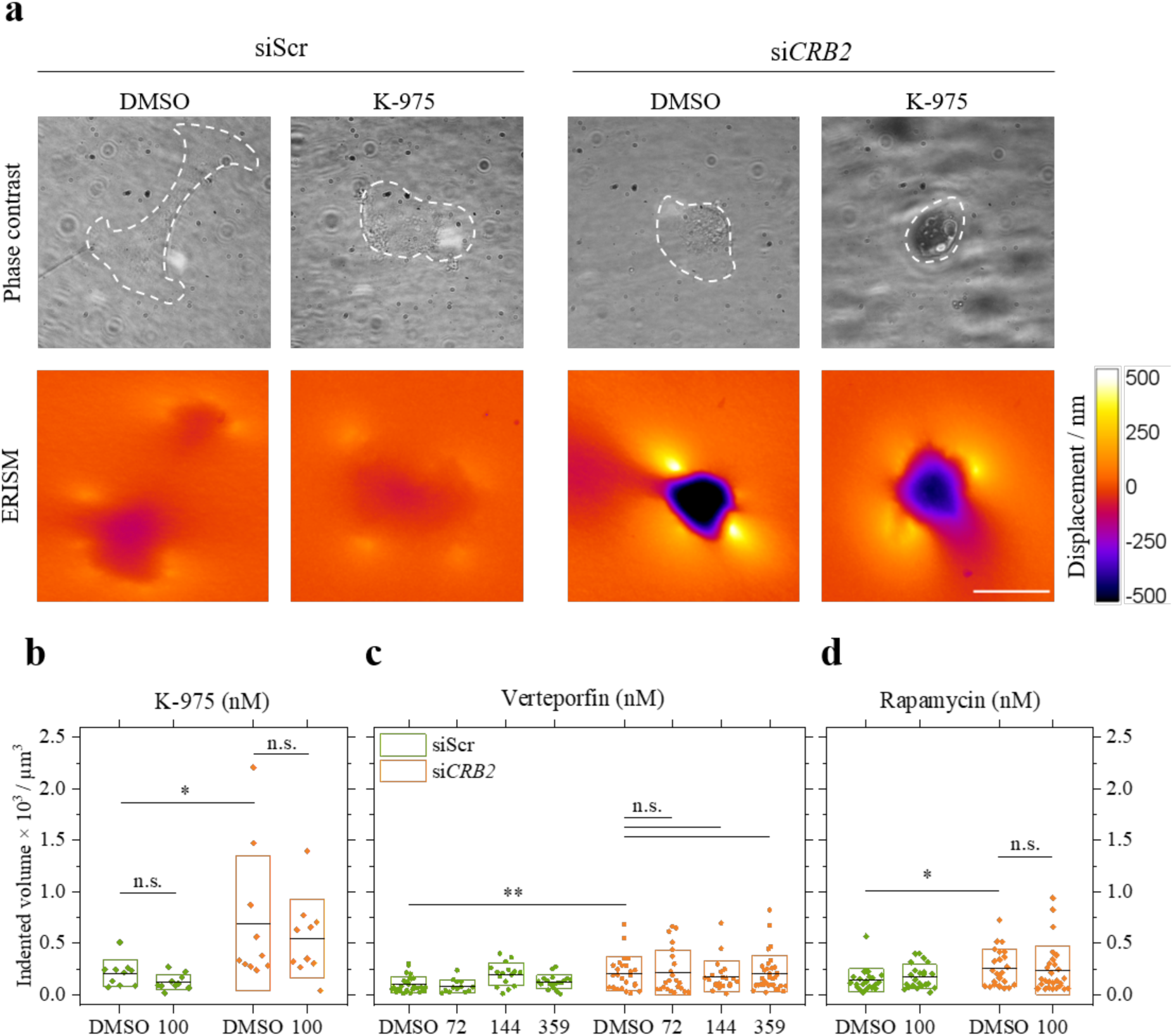
Hippo and mTOR signaling related drug treatments. **a)** Phase contrast images and ERISM displacement maps of representative differentiated control (siScr) and *CRB2* knockdown (si*CRB2*) podocytes on an ERISM substrate with an apparent stiffness of 53 kPa. Cell outlines are indicated by white-dashed lines. Groups were treated without (DMSO) or with 100 nM of K-975 for 24 hours. **b – d)** Mean indented volume (symbols), means (central lines) and ±SD (boxes) of single podocytes after 24 h of treatment with **b)** 100 nM of K-975, **c)** verteporfin with various of concentrations, and **d)** 100 nM of rapamycin for siScr and si*CRB2* podocytes. All groups were treated with 0.2% of DMSO. *N* ≥ 10. Scale bar, 50 μm. Groups were compared using two-tailed *t*-tests with non-equal variance, n.s.: no significance, *: *p* ≤ 0.05, **: *p* ≤ 0.01, ***: *p* ≤ 0.001.

The findings of the experiments with YAP-inhibitors suggest that the increase in contractility of CRB2 knockdown cells may not be exclusively a result of YAP activation, but that other pathways may also play a role or that cells can activate compensation pathways if YAP is inhibited via drugs.

## DISCUSSION

In this study, we report the identification of a rare compound heterozygous mutation in *Crumbs homolog 2* (*CRB2*; *FSGS9*) in an East Asian kindred with early-onset FSGS/SRNS. To model the effects of the disease-causing compound heterozygous mutation in Family DUK40595, we established stably-expressing scrambled (siScr) and *CRB2* (si*CRB2*) siRNA lines to determine if CRB2 functions upstream of YAP regulate podocyte mechanotransduction signaling. We showed that CRB2 knockdown significantly enhanced YAP/TEAD reporter activity, YAP target gene expression, and mechanotranductory signaling. CRB2 knockdown also significantly reduces podocyte cell area, upregulates F-actin and mechanosensitive protein expression and increases focal adhesion (FA) density. Additionally, we employed ERISM at various substrate stiffnesses to measure the effects of CRB2 knockdown on podocyte force development during differentiation. siScr and si*CRB2* podocytes demonstrated contractile forces transmitted at FA sites that were primarily distributed along the cell periphery. We found that CRB2 knockdown increases podocyte contractility in a substrate stiffness-dependent manner. The knockdown effect decreased with increasing substrate stiffness, indicating impaired mechanosensing in CRB2 knockdown cells at low substrate stiffness.

Together, these results strongly suggest a role of CRB2 as upstream activator of YAP activity. In siScr cells, we hypothesize that YAP is deactivated by phosphorylation through CRB2-MST1/2-LATS-mediated Hippo pathway signaling (Figure 8a). This posttranslational modification prevents YAP from entering the nucleus and can lead to its cytoplasmic sequestration with 14-3-3 protein or degradation (Figure 8a). In contrast, CRB2 knockdown may promote the downregulation of Hippo pathway signaling in podocytes, leading to an inappropriate upregulation of YAP activity and target gene expression (Figure 8b). Inappropriate upregulation of gene targets such as *THBS-1* could then lead to hyperactivation of integrin signaling, FAK activation, and FA growth (87) (Figure 8b). Increased YAP activity is also known to activate TGF-β/RhoA/ROCK signaling and myosin II activity (88). Both activation pathways could promote an increase in cell contractility (89, 90) (Figure 8b). Mechanical compression of the nucleus as a result of the increased cell contractility has been shown to directly lead to further nuclear translocation of YAP (91). Moreover, RhoA (via actomyosin) and FAK activity are also known to inhibit LATS1/2, which can lead to a further activation of YAP (92).

**Figure 8.**
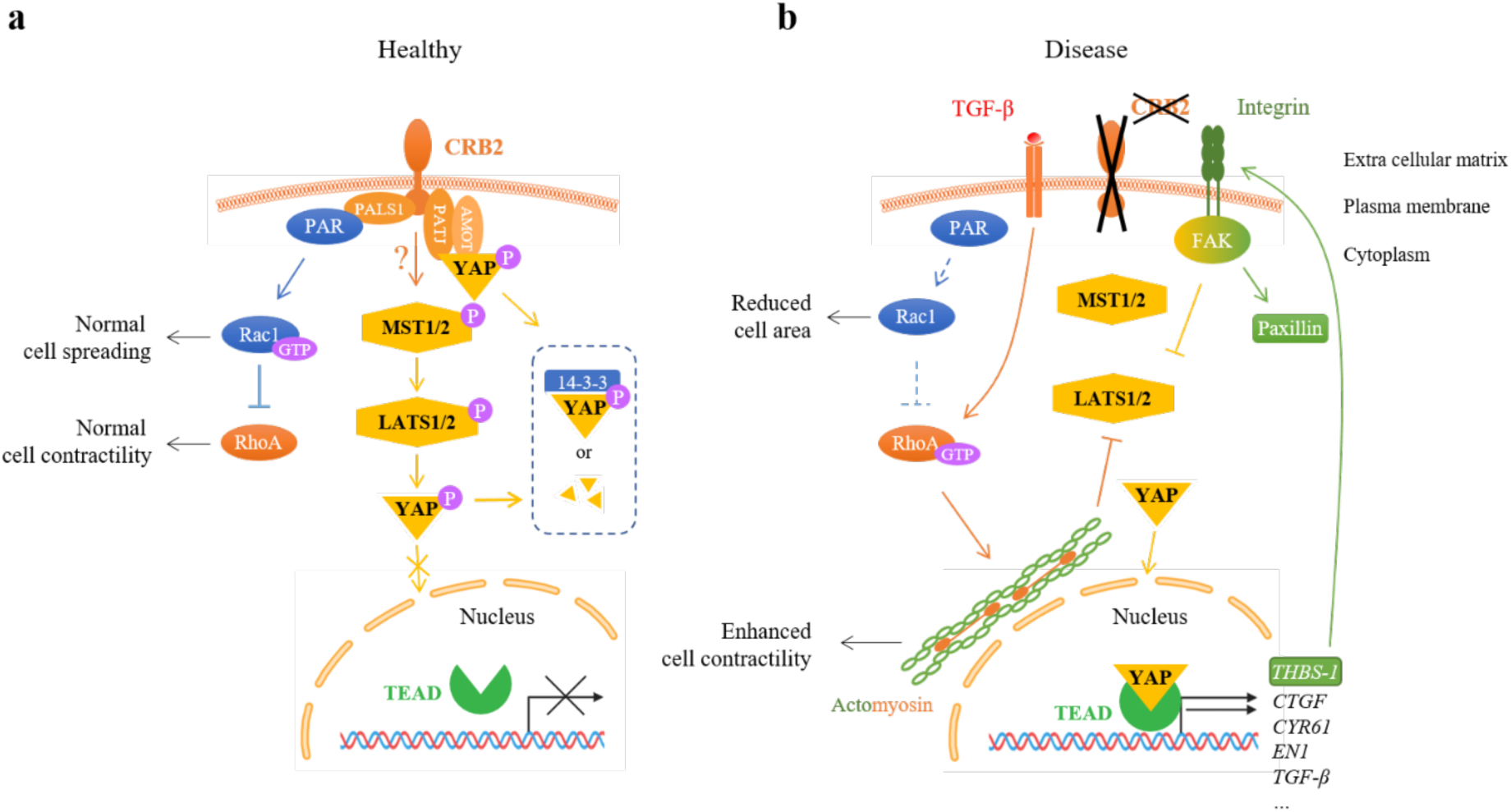
Proposed Effect of CRB2 Deficiency on Podocyte Mechanotransduction. Schematic diagram of the possible signaling pathways involved upon *CRB2* knockdown. **a)** In healthy podocytes, the Crumbs complex associates with the PAR complex, thereby facilitating normal cell spreading and contractility. Either the interaction with crumbs complex or the activation of MST1/2 and LATS1/2 signaling of the Hippo pathway may promote YAP phosphorylation (deactivation), leading to retention or degradation of cytoplasmic YAP. **b)** In *CRB2* mutant-related nephropathy, the activity of Rac1 might be suppressed by the disassociation of the PAR complex from the Crumbs complex. Together with the activated TGF-β signaling, this may lead to a higher activity of RhoA/ROCK/Myosin II signaling, thereby resulting in a smaller cell spreading area and an enhanced cell contractility. In the absence of CRB2, upregulated RhoA-and FAK-mediated LATS1/2 suppression may also result in nuclear translocation of YAP and upregulation of YAP target genes such as *THBS-1*, *CTGF*, *CYR61*, *EN1* and *TGF-β*, etc. FAK signaling is activated by THBS-1, leading to the upregulation of paxillin. Question marks, assumptions. Solid lines, normal signaling. Dashed lines, reduced signaling. AMOT, angiomotin. CRB2, crumbs 2. CTGF, connective tissue growth factor. CYR61, cysteine-rich angiogenic inducer 61. FAK, focal adhesion kinase. GTP, guanine triphosphate. LATS1/2, large tumor suppressor kinase 1/2. MST1/2, mammalian sterile 20-like kinase 1/2. P, phosphate. PAR, protease activated receptor complex. Rac1, Ras-related C3 botulinum toxin substrate 1. RhoA, Ras homolog family member A. ROCK, Rho-associated protein kinase. TEAD, transcriptional enhanced associate domain. TGF-β, transforming growth factor beta. THBS-1, thrombospondin 1. YAP, yes-associated protein.

Interestingly, we did not find evidence that the increased cell contractility of CRB2 knockdown cells was significantly reduced when cells were treated with the YAP inhibitors K-975 and verteporfin (84, 93). This might suggest that the increase in contractility of CRB2 knockdown cells is not only a direct result of YAP activation, but that other pathways also play a role. A possible alternative activation path is that the association of the PAR complex to PALS1 in the crumbs complex is disrupted upon CRB2 knockdown (94–96) (Figure 8). This might lead to reduced activation of Rac1 downstream of the PAR complex, resulting in the observed reduction in cell area (97), as well as to increased RhoA activity and thus the observed increase in cell contractility (81, 98).

It is important to point out that K-975 has not been applied in podocytes before. Based on prior work by Kaneda *et al.*, we selected a treatment dose of 100 nM, which was sufficient to reduce YAP target gene expression in multiple mesothelioma cell lines. Yet, in *in vitro* studies, dosing ranges up to 10 µM were evaluated with similar effects on gene expression (99). Similarly, with verteporfin, prior studies of the molecule in immortalized podocytes saw effective inhibition of YAP activity and gene expression with doses used as high as 5 µg/mL (100). However, in our study, doses higher than 1.0 µg/mL (1.5 µM) significantly reduced cell viability for both cell lines. With respect to rapamycin, the persistent activation of YAP in si*CRB2* podocytes may have limited the possibility of autophagic degradation in the absence of regular nuclear-to-cytoplasmic shuttling.

Significant progress has been made over the past decade in our understanding of disease-causing *CRB2* mutations in SRNS/FSGS. The first description by Ebrasi *et al.* clearly established *CRB2* mutations as a cause of autosomal recessive podocyte injury and impaired glomerular integrity (23). Moreover, using a *crb2*-deficient zebrafish model, they demonstrated that crb2 loss impaired apical basal polarity of podocytes, disrupted foot process morphology and arborization, altered trafficking of SD proteins such as nephrin, podocalyxin and ZO-1, disrupted slit diaphragm formation and compromised glomerular filtration barrier integrity (23). Since then, several studies have confirmed and expanded upon these findings documenting ∼30 human disease-causing *CRB2* mutations (27, 31, 32). Nonetheless, insight into the pathogenic mechanisms of CRB2-mediated podocyte injury remain limited. A recent report by Yang *et al.* suggested that novel pathogenic *CRB2* mutations reduce sphingosine-1-phosphate receptor 1 (S1PR1) expression/phosphorylation and that S1PR1-deficiency disrupts SD protein expression (32). This mechanism is plausible as S1PR1 signaling has been shown to be cytoprotective against podocyte injury in diabetic nephropathy, and disruption of this protection mechanism could predispose podocytes to dysfunction and loss (101, 102). Prior studies by Möller-Kerutt *et al.* showed that CRB2 is an essential SD protein that organizes in adjacent clusters with nephrin and that pathogenic mutations in *CRB2* may promote defective posttranslational modification and inappropriate retention in the endoplasmic reticulum (ER) leading to podocyte injury or loss by ER stress (26, 30). Additionally, Hamano *et al.* demonstrated that CRB2 associates with mTORC1 in developing podocytes and may influence mTORC1-mediated management of cellular energy resources (103). The changes in YAP activity in CRB2 knockdown podocytes presented by our data may be linked to Hippo pathway signaling. Prior studies of Hippo signaling in podocytes have shown that YAP activity is cytoprotective and that nuclear exclusion of the transcriptional co-regulator drives podocyte injury (37–39, 104, 105). Because YAP is recognized as a universal mechanotransducer (106), we focused on understanding the potential contribution of CRB2 to regulation of YAP-mediated mechanosignaling in podocytes. A prior study by Rinschen *et al.* showed that puromycin aminonucleoside (PAN)-induced glomerular injury in rats increased YAP activity and target gene expression (i.e. Cyr61/CCN1, CTGF/CCN2 and Ankrd1) and that inhibition of Hippo signaling with verteporfin reduced YAP activation and proteinuria. Additionally, they showed that podocyte YAP activity was influenced by substrate rigidity and F-actin (42). Although the Rinschen study did not address the role of CRB2, our findings related to YAP activation and mechanosignaling in podocytes were consistent.

While the majority of our results suggests a link between CRB2 and YAP activity, our data does not prove whether this link is made through Hippo pathway signaling, e.g. via the activation of MST1/2 and LATS1/2 (Figure 8a). It has been reported in the literature that YAP also directly associates with the crumbs complex (Figure 8a). This association could potentially promote cytoplasmic retention/degradation of YAP without the involvement of the Hippo pathway (107).

In conclusion, we identified a novel human CRB2 gene variant contributing to familial FSGS in combination with a known variant. CRB2 knockdown in human urinary podocytes resulted in increased YAP and TEAD gene expression, enhanced YAP and YAP-target protein levels, and higher cell contractility. In addition, CRB2 knockdown reduces the ability of podocytes to adapt their contractile force to the stiffness of their substrate (mechanotransduction). We also noted elevated F-actin, paxillin, and FAK protein expression induced in CRB2 deficient podocytes. Our results suggest that CRB2 is an upstream regulator of YAP-mediated mechanotransduction signaling in podocytes, even though other pathways may also play a role in mechanosignaling downstream of CRB2. These findings provide crucial mechanical insights for understanding *CRB2* mutation-induced podocytopathy.

## DATA AVAILABILITY

The data sets supporting this publication can be accessed via the PURE repository of the University of St Andrews at <DOI to be assigned upon acceptance>.

## SUPPLEMENTAL MATERIAL

Supplementary Figures S1 – S4 accompany this submission.

## ACKNOWLEDGEMENTS

The authors thank Dr. Rasheed A. Gbadegesin and Megan Chryst-Stangl of the Duke Genetics of Kidney Disease Study and the families of the Duke FSGS Study. Viral Vectors were provided by Duke University Viral (https://sites.duke.edu/dvvc/). Immunoassays were performed in the Biomarkers Core Facility at the Duke Molecular Physiology Institute.

## DISCLAIMERS

None.

## AUTHOR CONTRIBUTIONS

M.C.G, N.M.K. and G.H. conceived and designed research; G.H., S.S., R.R., C.J.S., and S.I. secured patient samples and performed diagnostic analyses; Y.S., S.S., M.E.K., B.S., S.I., X.T. and S.N.D. performed experiments; Y.S., N.M.K. and G.H. analyzed data; Y.S., N.M.K., M.C.G. and G.H. interpreted results of experiments; Y.S., N.M.K. and G.H. prepared figures, Y.S. drafted manuscript, All authors contributed substantively to the editing and revised manuscript, M.C.G. and G.H. approved the final version of the manuscript.

---

**Figure S1:**
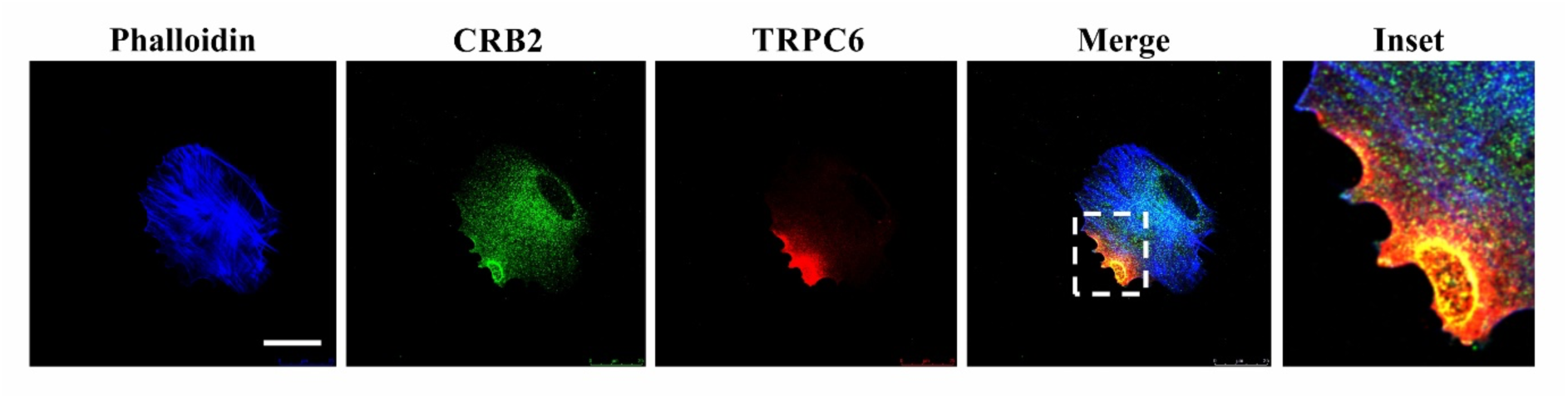
*CRB2* colocalizes with TRPC6 in immortalized human podocytes. *CRB2* is expressed throughout the apical membrane of mature, immortalized human podocytes and demonstrates regional colocalization with the slit diaphragm protein TRPC6 (Inset). Scale bar, 25 µm.

**Figure S2:**
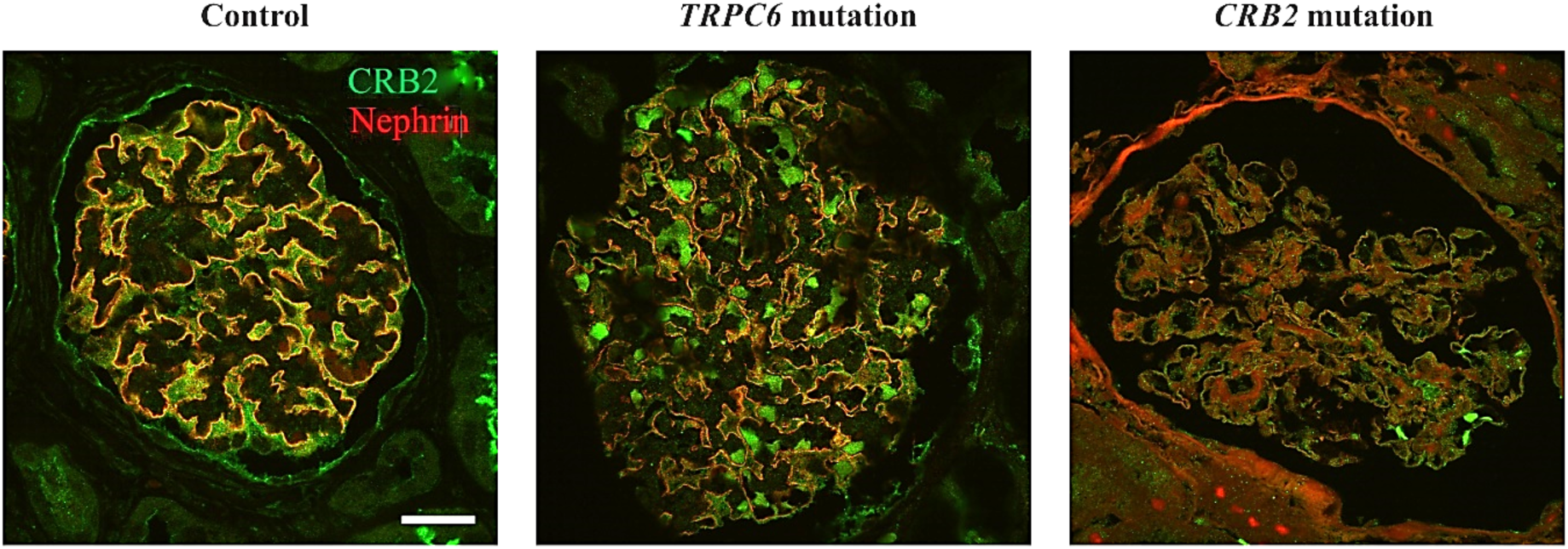
*CRB2* staining in glomeruli bearing the p.Gly1036Alafs*43 truncating mutation. Representative kidney biopsy images with *CRB2* and Nephrin staining in a healthy human glomerulus (control), a patient with FSGS due to a pathogenic *TRPC6* mutation and a patient with biopsy proven FSGS bearing the pathogenic truncating p.Gly1036Alafs*43 mutation. The uniform distribution of podocyte *CRB2* staining in the control is largely preserved but mildly irregular in the patient with the *TRPC6* mutation. Notably, in the patient with FSGS due to the truncating *CRB2* mutation, *CRB2* staining is obliterated, consistent with the predicted effect of the mutation to prevent the transmembrane insertion of the protein.

**Figure S3:**
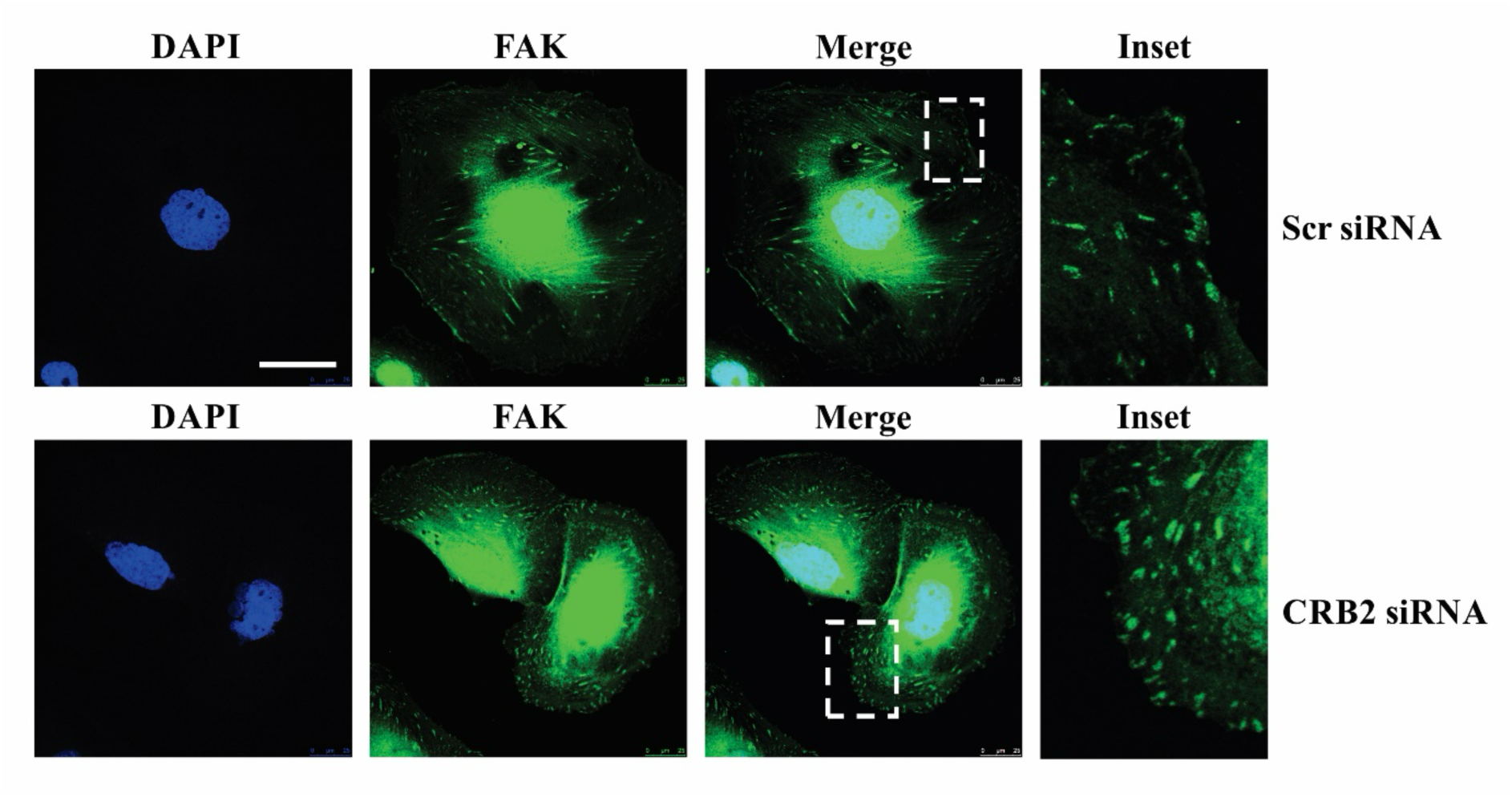
Focal adhesion density is enhanced in *CRB2* KD podocytes. Focal Adhesion Kinase (FAK) staining is used to identify focal adhesions by indirect immunofluorescence imaging in siScr and si*CRB2* podocytes (Insets). Although the number of focal adhesions per cell is similar in both lines, the focal adhesion density is greater in si*CRB2* podocytes due to their significantly reduced cell area as shown in Figure 2c of the main text. Scale bar, 25 µm.

**Figure S4:**
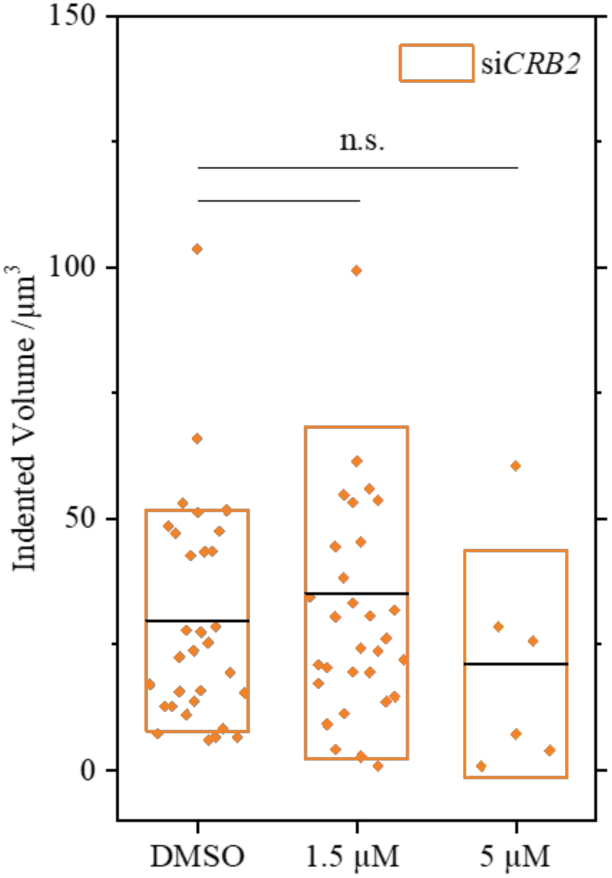
No reduction in cell contractility of *CRB2* KD podocytes upon treatment with verteporfin at high dose. Mean indented volume (symbols), means (central lines) and ±SD (boxes) of individual 12-day differentiated podocytes after 24 h of verteporfin treatment at two different concentrations in dark and in DMSO control, for *CRB2* knockdown (si*CRB2*) podocytes. Excessive cell death was observed for si*CRB2* podocytes treated with 5 µM of verteporfin. All groups were treated with 0.2% of DMSO and *N ≥* 6. Groups were compared using two-tailed *t*-tests with non-equal variance, n.s.: no significance, *: *p* ≤ 0.05, **: *p* ≤ 0.01, ***: *p* ≤ 0.001.

